# From dimer to tetramer: the evolutionary trajectory of C4 photosynthetic-NADP-ME oligomeric state in Poaceae

**DOI:** 10.1101/2025.01.05.631420

**Authors:** Jonas M. Böhm, Simone Willms, Oja Ferrao, Martin Buitrago-Arango, Meike Hüdig, Gereon Poschmann, Nazanin Fazelnia, Luitgard Nagel-Steger, Sebastián Klinke, Athina Drakonaki, Christos Gatsogiannis, Marcos A. Tronconi, Clarisa E. Alvarez, Veronica G. Maurino

## Abstract

The C4 carbon concentrating mechanism relies on specialized enzymes that have evolved unique expression patterns and biochemical properties distinct to their ancestral housekeeping forms. In maize and sorghum, the evolution of C4-NADP-malic enzyme (C4-NADP-ME) involved gene duplication and neofunctionalization, leading to the emergence of two plastidic isoforms: C4-NADP-ME and nonC4-NADP-ME, each with distinct kinetic and structural features. While C4-NADP-ME functions primarily as a tetramer, nonC4-NADP-ME exists in an equilibrium between dimeric and tetrameric forms, favoring the dimer in solution. This study shows which evolutionary changes in amino acid sequences influence the structure and function of these isoforms. By integrating X-ray crystallography, cryo-electron microscopy, computational molecular modeling and targeted biochemical analysis of mutant and truncated protein variants, we identify crucial roles for the N- and C-terminal regions and specific amino acid residues in governing isoform oligomerization. Our results reveal that the N-terminal region is essential for stabilizing the dimeric form of nonC4-NADP-ME, whereas specific adaptive substitutions and interactions with the C-terminal region enhance the stability of the tetrameric state characteristic of the C4-adapted isoform. We propose that differences in the N-terminal domain between the C4 and nonC4 isoforms reflect distinct selective pressures, which have driven their evolutionary divergence to fulfill specialized cellular functions.

## Introduction

The evolution of photosynthesis in land plants reveals a fascinating adaptation: the C4 carbon concentrating mechanism (Sage *et al*., 2012). This remarkable metabolic pathway enhances photosynthetic efficiency by concentrating CO_2_ around Ribulose-1,5-bisphosphate carboxylase-oxygenase (Rubisco) by reducing its wasteful oxygen fixation reaction. The C4 pathway has independently evolved in at least 45 different lineages across 19 families of flowering plants – a striking example of convergent evolution (Sage *et al*., 2011). In most C4 lineages, NADP-malic enzyme (C4-NADP-ME), specifically localized in the chloroplasts of bundle sheath cells, catalyzes the decarboxylation of malate, thereby supplying a concentrated source of CO₂ for Rubisco (Furbank, 2011). This specialized C4-NADP-ME evolved from a plastidic housekeeping isoform through gene duplication and subsequent modifications in the promoter and 5′ region, enabling high expression levels in bundle sheath cells (BSCs), and further protein neo-functionalization (Tausta *et al*., 2002, Christin *et al*., 2009, Brown *et al*., 2011, Alvarez *et al*., 2019).

The NADP-ME gene family is diverse, with isoforms found in both the cytosol and plastids (Maurino *et al*., 2009, Tronconi *et al*., 2018). In addition to its role in C4 photosynthesis, non-photosynthetic isoforms of NADP-ME in the cytosol and plastids participate in other important processes, such as defense responses (Maurino *et al*., 2001, Chi *et al*., 2004, Voll *et al*., 2012), seed germination (Yazdanpanah *et al*., 2018), and lipid biosynthesis (Lai *et al*., 2002).

Amino acid substitutions that enabled C4-NADP-ME to participate in the C4 pathway are specific to certain genetically close C4 lineages (Alvarez *et al*., 2019, Alvarez and Maurino, 2023, Eckardt *et al*., 2024). In C4 plants like maize (*Zea mays*) and sorghum (*Sorghum bicolor*), members of the same C4 lineage (Andropogoneae) within the subfamily Panicoideae, the plastidic C4-NADP-ME and housekeeping (nonC4-NADP-ME) isoforms bear significant kinetic and structural differences. Compared to the plastidic nonC4 isoform, C4-NADP-ME in these species exhibits higher affinity for the substrate malate, an increased catalytic rate, and malate inhibition at pH 7.0 (Maurino *et al*., 2001, Maier *et al*., 2011). The pH-dependent malate inhibition was proposed to be important for the efficient operation of the C4 pathway by preventing CO_2_ loss and preserving the malate pool during the night when stromal pH drops due to the inactivity of the photosynthetic electron transport chain (Su and Lai, 2017, Alvarez *et al*., 2019, Bovdilova *et al*., 2019).

In maize and sorghum, the oligomeric state of C4-NADP-ME is also different from the nonC4 isoform. C4-NADP-ME forms a stable tetramer of around 250 kDa within the physiological chloroplast pH range of 7.0-8.0 (Alvarez *et al*., 2019, Bovdilova *et al*., 2019), whereas nonC4-NADP-ME predominantly exists as a dimer of 130 kDa, as observed through native PAGE (Saigo *et al*., 2013, Alvarez *et al*., 2019). In most organisms, NADP-ME exists in a predominant oligomeric state, typically forming dimers or tetramers with strong interactions at the dimer interface and weaker interactions at the tetramer interface (Xu *et al*., 1999, Yang *et al*., 2002a, Yang *et al*., 2002b, Alvarez *et al*., 2019, Grell *et al*., 2022). However, transitions between monomers, dimers and tetramers were observed in response to pH changes in the case of the pigeon liver cytosolic NADP-ME, indicating a dynamic equilibrium between the different oligomerization states (Chang *et al*., 1988). Similarly, isolated C4-NADP-ME from sugar cane (*Saccharum officinarum*) was found preferentially as dimers at pH 7.0 and as a tetramer at pH 8.0. Due to this observation, it has been hypothesized that C4-NADP-ME may alter its oligomeric state in response to changes in stromal pH, as a mechanism to preserve the malate pool during the night (Iglesias and Andreo, 1990).

Different structural elements of the monomers have been identified as key determinants of the oligomerization state and as critical factors driving the functional divergence of the NADP-ME isoforms. In human mitochondrial NADP-ME, the C-terminal loop interacts with the neighboring dimer, promoting tetramer formation (Grell *et al*., 2022). Additionally, in the other human mitochondrial isoform, NAD(P)-ME, a C-terminal extension of seven residues stabilizes the tetramer by interacting with both monomers of the opposing dimer (Xu *et al*., 1999). However, bacterial, yeast, and plant NADP-MEs lack this C-terminal extension, suggesting that alternative structural mechanisms govern their oligomeric organization. Plant NADP-MEs possess a longer N-terminal region, although of variable length, than that of human enzymes, and recent studies have linked this region to changes in their quaternary structure. For instance, the deletion of the first 20 amino acid residues from the mature dimeric maize nonC4-NADP-ME induces tetramer formation (Saigo *et al*., 2013, Alvarez *et al*., 2019). Interestingly, although the transit peptide of maize and sorghum plastidic NADP-ME isoforms originated from a single evolutionary event (Tausta *et al*., 2002, Christin *et al*., 2009), the C4 isoform contains a shorter N-terminal region due to a 15-residues deletion (Alvarez *et al*., 2019). Despite these insights, the precise function of the N-terminal region in regulating oligomerization remains unclear, partly due to its unresolved structure in crystallized C4-NADP-MEs (Alvarez *et al*., 2019).

Beyond the N-terminal differences, previous work has identified 20 residues as potentially critical to the evolution of maize and sorghum C4-NADP-ME, based on their differential conservation between the C4- and nonC4 isoforms (Alvarez *et al*., 2019). A strictly differentially substituted position is one where all C4 isoforms share the same amino acid, while housekeeping isoforms from C4 and C3 plants share a different amino acid (Alvarez *et al*., 2019, Hüdig *et al*., 2022). Four of these residues have previously been shown to contribute to the specific C4 characteristics of maize C4-NADP-ME, and can therefore be considered adaptive substitutions (Alvarez *et al*., 2019).

In this study, we investigate the structural elements underlying the oligomerization differences between the C4- and nonC4-NADP-ME isoforms. Specifically, we focus on the role of the N-terminal region and the differentially conserved amino acids in driving the structural divergence of the isoforms in maize and sorghum. Through biochemical and structural analyses of recombinant wild-type NADP-ME isoforms and their mutants, combined with predictive algorithms, we showed that mutations at the dimer interface and N-terminal modifications selectively influence the oligomeric states of plastidic housekeeping and C4-photosynthetic NADP-ME. Alongside *cis*-element analysis at 5’ coding regions, we provide insights into the evolutionary pathway leading to the differential oligomerization of these plastidic isoforms.

These results will help to clarify how these structural variations influence the specific function of NADP-ME isoforms, and may offer broader insights into the evolution of enzymes and their functional diversification in plants.

## Results

### Oligomeric organization of nonC4- and C4-NADP-ME in maize tissues

Our previous crystallographic analysis of recombinant C4-NADP-ME from maize and sorghum revealed that the protein is a tetramer formed by a dimer of dimers (Alvarez *et al*., 2019). Analytical ultracentrifugation (AUC) measurements conducted in this study at pH 8.0 (sedimentation coefficient of 10 S; Fig. 1A) along with previous analysis at pH 7.0 (Bovdilova *et al*., 2019), confirm the stable tetrameric state of recombinant maize C4-NADP-ME. In contrast, AUC analysis of nonC4-NADP-ME indicated that the enzyme is in a dynamic dimer-tetramer equilibrium in solution, with a notable tendency towards aggregation at both pH values (Fig. 1A and Suppl. Fig. 1). The AUC distribution profile for nonC4-NADP-ME at pH 8.0 shows variability in oligomeric states depending on the protein concentration, with higher concentration favoring higher order oligomers (sedimentation coefficient around 7.0 for the dimer; Fig. 1A). Consistent with this observation, under native PAGE conditions, tetrameric recombinant C4-NADP-ME migrates between the 445 and 146 kDa markers (Fig. 1B), whereas recombinant nonC4-NADP-ME generally shows both oligomeric states, with the highly predominant dimeric form migrating just below the 146 kDa marker (Fig. 1B). It is worth mentioning that generally only the dimeric state is observed after native PAGE.

**Figure 1.**
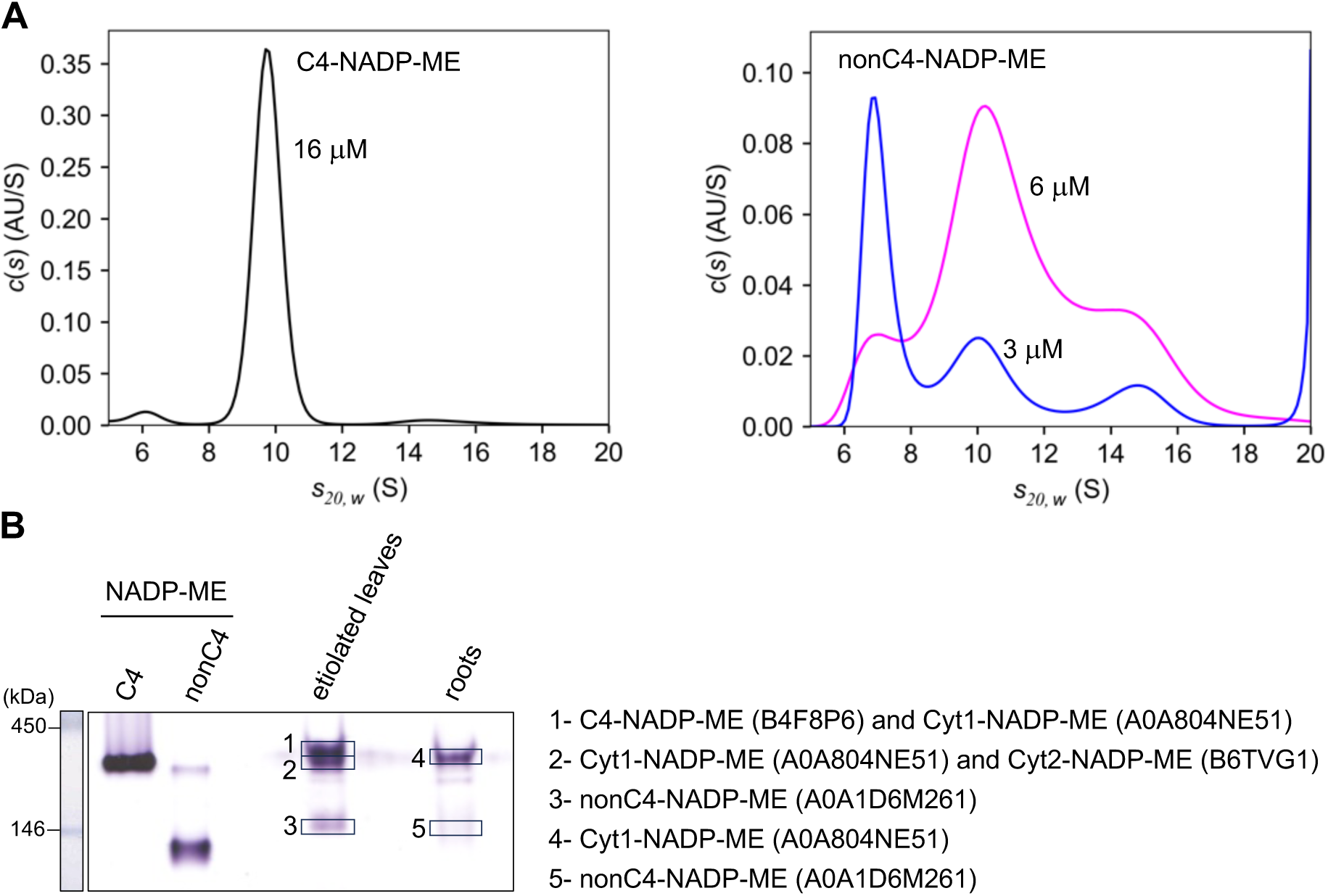
Analytical ultracentrifugation and native PAGE analysis. **A.** Continuous sedimentation coefficient distribution for maize C4- and nonC4-NADP-ME at pH 8.0. The distribution profile for nonC4-NADP-ME shows variability in oligomeric states (dimers and tetramers) depending on the protein concentration. The protein concentration analyzed are indicated in the graphics. Data were analyzed using the c(s) model in the software package SEDFIT. **B.** Native PAGE (7%) analysis of 35 µg of protein extract from etiolated leaf and root tissue coupled to an in-gel NADP-ME activity assay. The excised protein bands and the corresponding identified NADP-ME isoforms are shown on the right. Recombinant C4- and nonC4-NADP-ME were used as controls (0.5 µg) and SERVA native marker is shown on the left. Bands with NADP-ME activity were analyzed by MS and the identified NADP-ME isoforms are listed on the right.

To further investigate the differences in oligomeric states of C4- and nonC4-NADP-ME in plant tissues, we analyzed extracts of maize from etiolated leaves and roots using native PAGE coupled with an in-gel NADP-ME activity assay. All protein bands exhibiting NADP-ME activity were excised and analyzed by mass spectrometry. C4-NADP-ME and other two cytosolic isoforms (Cyt1-NADP-ME [Cyt1]; A0A804NE51 and Cyt2-NADP-ME [Cyt2]; B6TVG1) were identified in bands with mobilities similar to the recombinant C4-NADP-ME (Fig. 1B), indicating that they form tetramers. In both tissue types, the plastidic nonC4-NADP-ME appeared in a band with similar mobility compared to its recombinant counterpart, suggesting that it forms dimers (Fig. 1B). A slight difference in mobility was observed between the recombinant protein and the plant tissue-derived version, suggesting that, in plant tissues, nonC4-NADP-ME may adopt a more native conformation, interact with other proteins in the extract, or undergo post-translational modifications beyond the capabilities of the prokaryotic expression system.

Overall, our findings indicate that C4- and nonC4-NADP-ME exhibit differences in their oligomeric organization, both as recombinant proteins and following extraction from maize tissues.

### The N-terminal sequence of nonC4-NADP-ME is involved in its oligomeric organization

In previous work we showed that partial deletion of the N-terminal region of maize nonC4-NADP-ME, by eliminating the first 20 residues of the mature protein (nonC4DelN; Fig. 2A), displaces the oligomeric equilibrium of the protein to the formation of tetramers, suggesting that this region may be involved in the stabilization of the dimer, preventing tetramerization (Alvarez *et al*., 2019). Interestingly, there are notable differences between maize (and sorghum) nonC4-NADP-ME and C4-NADP-ME in the length of their N-terminal regions. The nonC4-NADP-ME isoform possesses a longer N-terminus, featuring a sequence of 15 residues (AAGVVVEDHYGEDSA) that was lost during the evolution of C4-NADP-ME (Fig. 2A). We hypothesize that this structural difference may have implications for the evolutionary adaptation of both enzymes.

**Figure 2.**
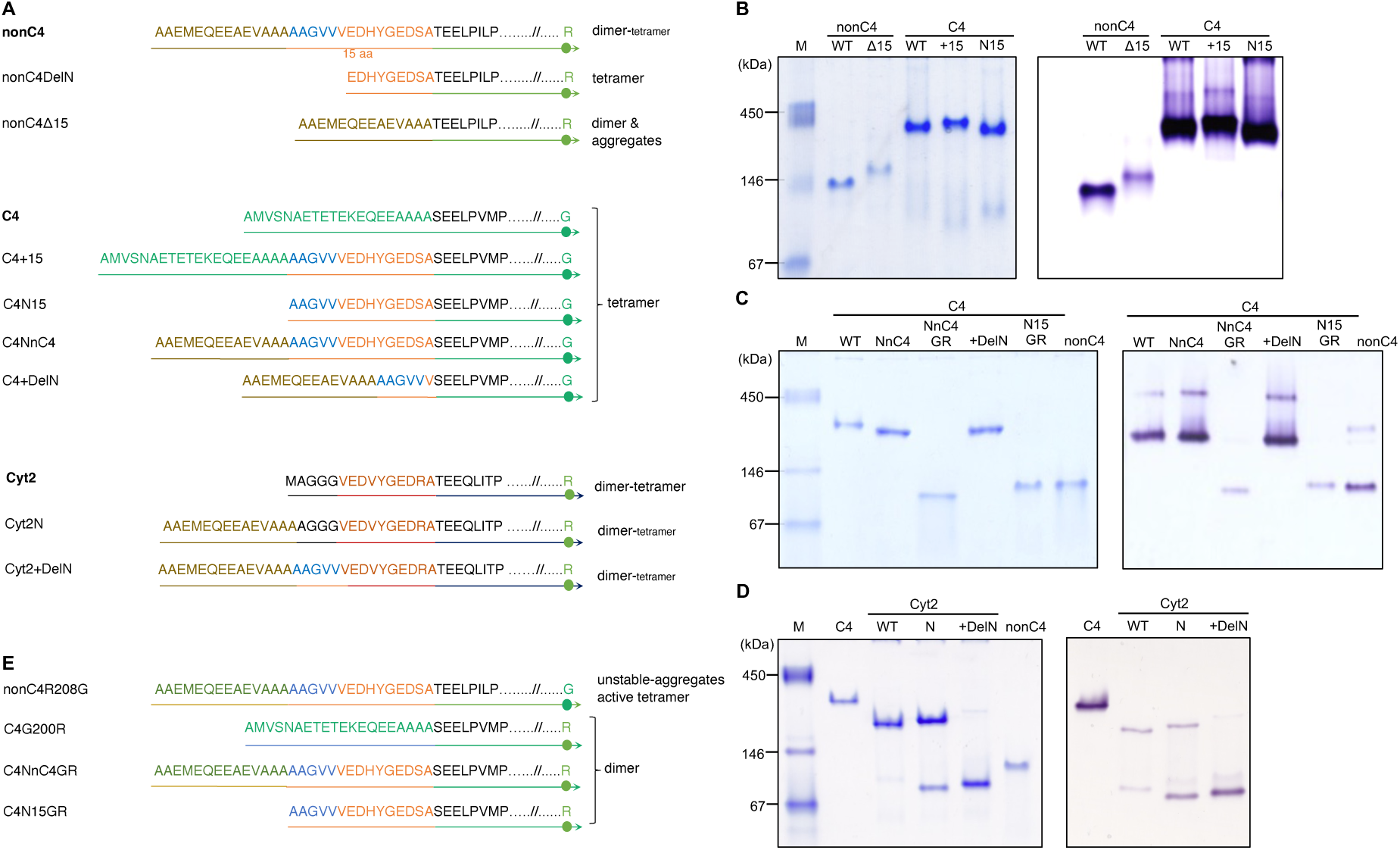
Analysis of C4- and nonC4-NADP-ME N-terminal variants. **A.** Overview of the chimeric and truncated N-terminal regions of the mature protein variants produced. **B.** Native PAGE (6%) of recombinant C4- and nonC4-NADP-ME variants with 5 μg protein loaded per lane followed by a Coomassie staining (left), or with 3 μg protein loaded per lane followed by an in-gel NADP-ME activity assay (right). **C-D.** Native PAGE (7%) of recombinant C4-NADP-ME (**C**) and Cyt2 (**D**) variants with 3 µg protein per lane followed by a Coomassie staining (left), or with 1 μg protein loaded per lane followed by an in-gel NADP-ME activity assay (right). In panels B-D, the SERVA native marker is shown on the left. **E.** Overview of the mutants produced at residue 200 in chimeric and truncated C4-NADP-ME variants and at the corresponding residue 208 in nonC4-NADP-ME.

In the nonC4DelN mutant, the first six (AAGVVV) of the 15 residues were deleted (Fig. 2A). Thus, to explore the role of the 15 amino acids present in nonC4-NADP-ME and their putative contribution to the differences in oligomeric organization between the isoforms, we generated several enzyme variants (Fig. 2A). Specifically, we created: (i) a nonC4-NADP-ME variant lacking the 15 residues (nonC4Δ15), (ii) a C4-NADP-ME variant with an extended N-terminal region by inserting the 15 residues from nonC4-NADP-ME at the position they are found in this isoform (C4+15); (iii) a C4-NADP-ME with only the 15 residues of nonC4 isoform at the N-terminal (C4N15); (iv) a C4-NADP-ME variant where the N-terminal region was replaced for the nonC4-NADP-ME N-terminal region - until the end of the 15 residues - (C4NnC4); and (v) a C4-NADP-ME variant with the N-terminal region of nonC4-NADP-ME until the end of nonC4DelN and thus lacking the EDHYGEDSA sequence (C4+DelN). These modified proteins were expressed in *Escherichia coli* and subsequently isolated using the immobilized metal affinity chromatography (IMAC) technology. Native PAGE coupled with in-gel activity assay was conducted to analyze the recombinant protein oligomeric states based on their migrations. We observed that all C4-NADP-ME variants presented similar mobilities to that of the wild-type enzyme, and they retained similar enzymatic activities (Fig. 2B and C). These results indicate that the N-terminal region of the C4-NADP-ME is not involved in the tetramerization of the protein.

The removal of the 15 residues from nonC4-NADP-ME (nonC4Δ15) resulted in a variant with slightly lower mobility in native PAGE compared to the original nonC4-NADP-ME isoform (Fig. 2B), likely due to changes in charge and/or conformation within the mutant protein. This modified protein also exhibits a tendency to aggregate, which may explain its challenging recovery in solution. Accordingly, AUC analysis of recombinant nonC4Δ15 revealed that the 15 amino acids deletion alters the aggregation behavior of the subunits, shifting them to higher oligomeric states, as compared to nonC4-NADP-ME (Suppl. Fig. 1). These results suggest that the 15 residues sequence AAGVVVEDHYGEDSA in the N-terminal region of nonC4-NADP-ME contains elements crucial for stable oligomeric assembly, as well as features that restrict tetramer formation. The nonC4DelN variant assembles as a tetramer (Alvarez *et al*., 2019), strongly suggesting that the amino acid sequence AAGVV plays a key role in this restriction.

To further investigate this, we expanded our analysis to a cytosolic NADP-ME from maize, whose ancestor is thought to have given rise to the plastidic isoforms by the acquisition of a transit peptide (Christin *et al*., 2009). To pinpoint the most likely ancestral cytosolic isoform, we conducted a phylogenetic analysis (Suppl. Fig. 2). This analysis incorporated 58 coding sequences from 16 grass species (Poaceae), along with sequences from *Joinvillea ascendens*, a species from a closely related sister family (Joinvilleaceae), and *Acorus americanus* (Acoraceae) as a reference for the last common ancestor of all monocots (Givnish *et al*., 2018). In the monocots, we identified four distinct, high-confidence NADP-ME clades (Suppl. Fig. 2). Notably, the plastidic NADP-ME clade forms a sister group to a set of cytosolic isoforms, strongly indicating that plastid-localized NADP-MEs evolved from these cytosolic ancestors. Our analysis supports the hypothesis that maize Cyt2 and nonC4-NADP-ME share a common evolutionary origin.

Comparative sequence analysis showed that Cyt2 has a shorter N-terminus than nonC4-NADP-ME, lacking the full N-terminal region of the nonC4 isoform until the end of the five-residue stretch AAGVV (Fig. 2A). Our analysis of extracts of maize from etiolated leaves and roots by native PAGE coupled with an in-gel NADP-ME activity assay showed that Cyt2 is present as tetramer (Fig. 1B). To investigate the structural and functional implications of the N-terminal differences between nonC4-NADP-ME and Cyt2, we expressed the wild-type Cyt2 and generated two N-terminal modified variants: Cyt2N, which includes only the N-terminal region of nonC4-NADP-ME without the AAGVV sequence, and Cyt2+DelN, which mirrors the N-terminal sequence of nonC4-NADP-ME including the AAGVV sequence (Fig. 2A). Native PAGE analysis of the recombinant variants revealed that wild-type Cyt2 predominantly exists as tetramer (Fig. 2D). Remarkably, introducing the N-terminal region of nonC4-NADP-ME without the AAGVV sequence shifted the equilibrium toward dimer formation. Furthermore, including the full N-terminal sequence with the AAGVV stretch induced a predominant dimer configuration, resembling the native oligomeric state of nonC4-NADP-ME.

Collectively, these findings provide strong evidence that in nonC4-NADP-ME the N-terminal extension facilitates dimer formation, with the AAGVV sequence serving as a key determinant in inhibiting the tetrameric assembly.

### The role of N-terminal modifications and adaptive substitutions in stabilizing tetramer formation in nonC4-NADP-ME

In a previous work we showed that residue F140 plays a crucial role in stabilizing the oligomeric structure of C4-NADP-ME (Alvarez *et al*., 2019). To explore this stabilizing effect in the N-terminal variants, we introduced a point mutation at F140 in C4-NADP-ME and the corresponding I148 in nonC4-NADP-ME N-terminal variants. The recombinant mutans C4-NADP-ME+15F140I (C4+15F140I), C4-NADP-MEN15F140I (C4N15F140I), and nonC4-NADP-MEΔ15148F (nonC4Δ15I48F; Δ15IF) were produced, isolated, and analyzed using native PAGE (Fig. 3A-B).

**Figure 3.**
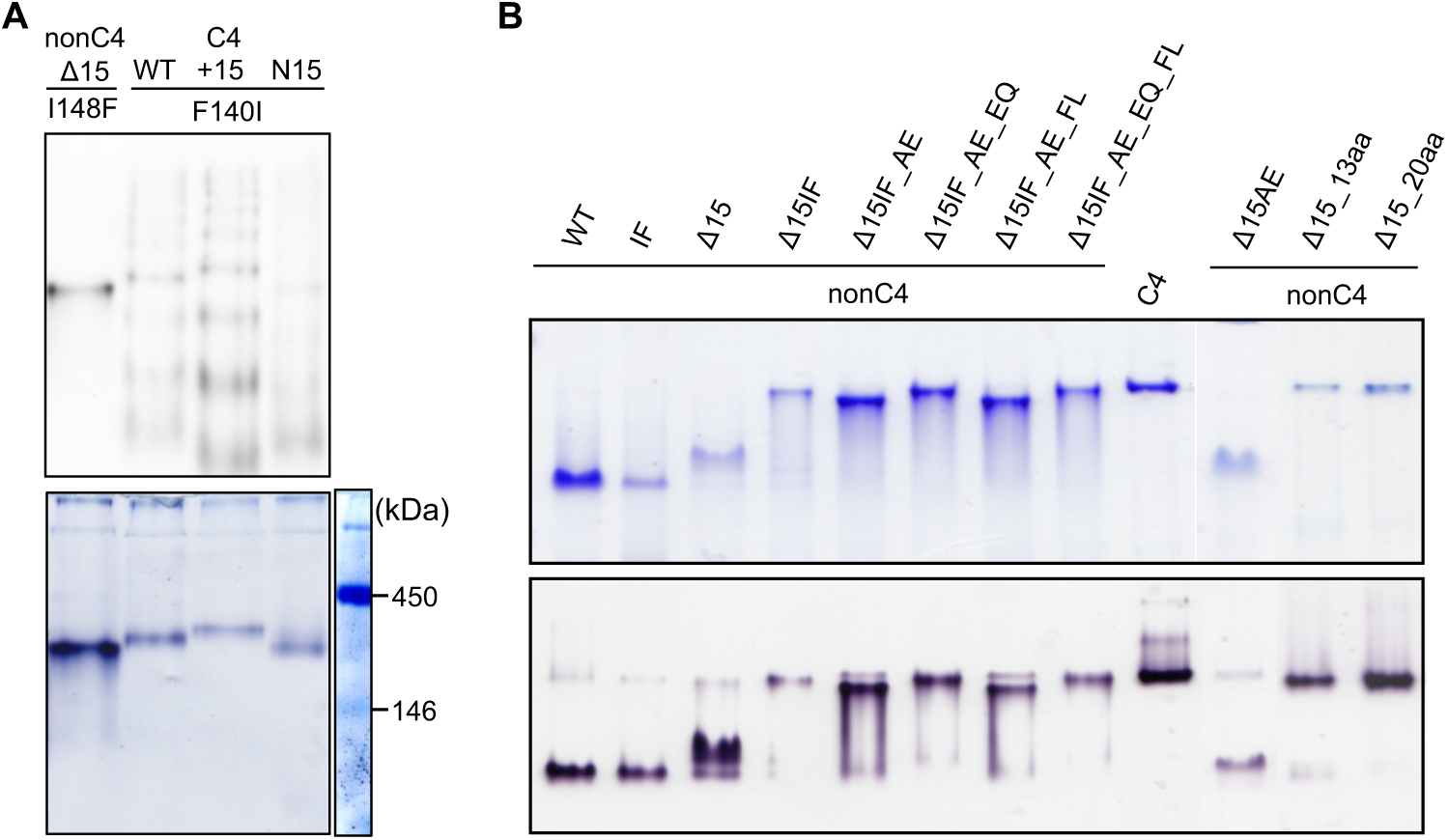
Native PAGE analysis of NADP-ME variants with combined N-terminal modifications and adaptive mutations. **A.** Native PAGE (7%) analysis of recombinant nonC4- and C4-NADP-ME N-terminal variants, each with a point mutation at residues 148 or 140, respectively. Top panel: Immunoblot with 5 μg protein per lane using anti-His-HRP antibodies. Bottom panel: In-gel NADP-ME activity assay with 3 μg protein per lane. The SERVA native marker is shown on the right. **B.** Native PAGE (7%) analysis of recombinant nonC4-NADP-ME variants. Top panel: Coomassie staining with 3 μg protein per lane. Bottom panel: In-gel NADP-ME activity assay with 1 μg of protein per lane. Due to low protein recovery, 2.5 μg of IF and 1.5 μg Δ15 and Δ15IF were loaded. Recombinant C4- and nonC4-NADP-ME were used as controls. IF, AE, EQ, and FL correspond to the mutations I148F, A347E, E511Q, and F552L in nonC4-NADP-ME. In the nonC4-NADP-ME variant where 13 residues were simultaneously mutated (13aa), the enzyme contains the following mutations: F100T, I148F, H150N, D167N, R171T, N172D, E185V, R208G, Q209R, R274D, A347E, E511Q, and F552L. In the nonC4-NADP-ME variant where 20 residues were simultaneously mutated (20aa), the enzyme contains the following mutations: F100T, I148F, H150N, D167N, R171T, N172D, E185V, R208G, Q209R, R274D, H312D, I325F, A347E, V377M, H385Q, V482I, E511Q, N514T, D529A, and F552L (Suppl. Table 1).

Our results show that all C4-NADP-ME variants carrying the F140I mutation exist as a mixture of oligomeric states and/or conformations (Fig. 3A). In-gel activity assays revealed that only the tetrameric forms of these variants are catalytically active (Fig. 3A). These findings undescore the importance of F140 in stabilizing the quaternary structure of maize C4-NADP-ME, regardless of N-terminal modifications.

Interestingly, the introduction of the I148F mutation in the nonC4Δ15 variant shifted the equilibrium from dimers toward active tetramer formation (Fig. 3B). Additionally, Δ15IF showed a tendency to aggregate, as indicated by AUC analysis (Suppl. Fig. 1), limiting the recovery of protein in solution. Notably, introducing the I148F mutation in nonC4-NADP-ME without the N-terminal deletion did not alter its oligomeric state (Fig. 3B), underscoring the crucial role of the N-terminal region in modulating oligomerization in this isoform.

Among the 20 amino acids differentially substituted between nonC4- and C4-NADP-ME in maize and sorghum, in addition to I148F, three substitutions, A347E, E511Q, and F552L (refer to Suppl. Table 1 for the positional homologous residue in C4- and nonC4-NADP-ME), are important for enhancing malate affinity in C4-NADP-ME (Alvarez *et al*., 2019, Eckardt *et al*., 2024). To further investigate this, we produced recombinant nonC4Δ15 proteins with combinations of these four mutations. All mutant proteins exhibited greater tetrameric stability compared to Δ15IF (Fig. 3B), although some still show aggregation in solution. Remarkably, the nonC4Δ15 variant containing a combination of a high number of adaptive substitutions (Δ15_13aa and Δ15_20aa) exhibited a highly stable tetrameric state, comparable with the wild-type C4-NADP-ME when all 20 amino acid substitutions were present (Fig. 3B). These findings indicate that the N-terminal region is crucial for maintaining the dimeric form of wild-type nonC4-NADP-ME. However, in the absence of the N-terminal region, the introduction of adaptive mutations plays a key role in stabilizing the tetrameric state of nonC4-NADP-ME.

### Identification of residues involved in the quaternary state shift from nonC4- to C4-NADP-ME

Previous analysis of four out of the 20 differentially conserved amino acids identified F140 as a key contributor to the structural stability of C4-NADP-ME (Alvarez, 2019). Additionally, introducing multiple mutations of these differentially conserved amino acids into Δ15IF resulted in the formation of a stable tetramer (Fig. 3B). These findings suggest that residues beyond F140 may also contribute to the tetramerization of C4-NADP-ME. To further investigate this, we extended our analysis to assess whether any of the remaining 16 differentially conserved residues contribute to the quaternary structure differences between the C4 and nonC4 isoforms of NADP-ME. We produced mutants of C4-NADP-ME by individually substituting 16 residues with their corresponding counterparts from nonC4-NADP-ME (T92F, N142H, N159D, T163R, D164N, V177E, G200R, R201Q, D266R, D304H, F317I, M369V, Q377H, I474V, T506N and, A521D; refer to Suppl. Table 1 for the corresponding amino acids in C4- and nonC4-NADP-ME) through gene synthesis or site-directed mutagenesis, and the recombinant proteins were isolated using IMAC.

Of the 16 mutants of C4-NADP-ME, 13 retained a tetrameric organization and exhibited activity similar to the wild-type enzyme (Fig. 4A). However, three recombinant mutants (C4T163R, C4D164N, and C4G200R) showed changes in oligomerization in comparison to the wild type enzyme (Fig. 4B). Both C4T163R and C4D164N showed an equilibrium of dimers and tetramers. This was further confirmed by AUC, where C4D164N was found to be 48% dimeric and 34% tetrameric (Fig. 4C). Notably, the C4G200R mutant presented mobility similar to nonC4-NADP-ME in native PAGE, existing as a stable dimer (Fig. 4B), as confirmed by AUC (Fig. 4C).

**Figure 4.**
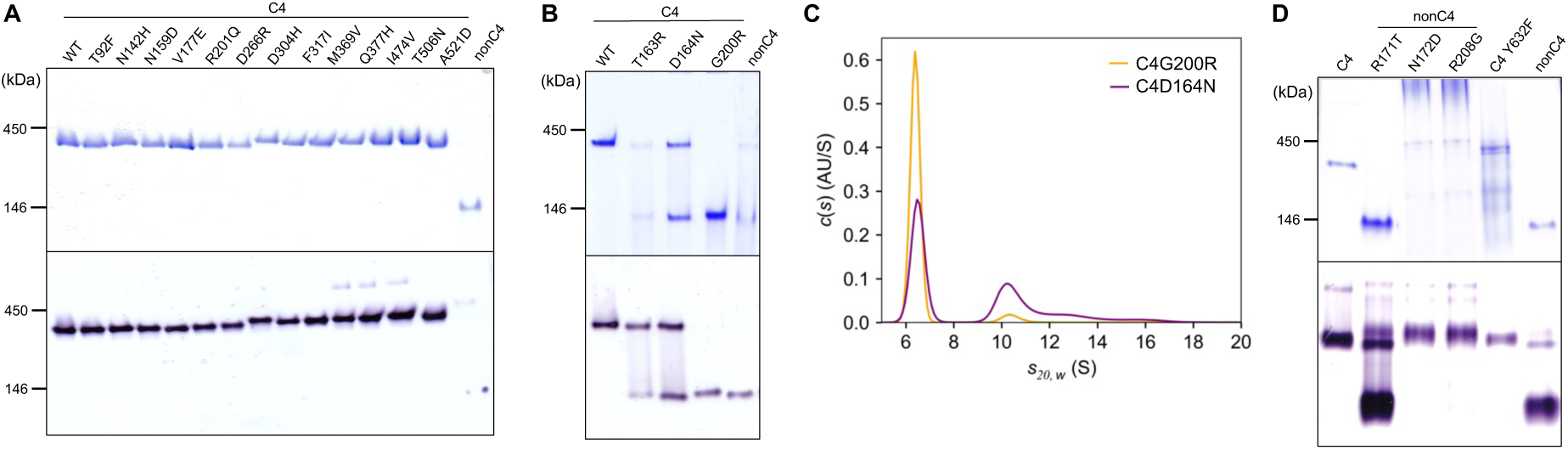
Native PAGE and AUC analysis of C4- and nonC4-NADP-ME variants with adaptive mutations. **A-B.** Native PAGE (7%) analysis of recombinant C4-NADP-ME variants. Top panel: Coomassie staining with 3 μg protein per lane. Bottom panel: *in-gel* NADP-ME activity assay with 1 μg protein per lane. **C.** Continuous sedimentation coefficient distribution of C4G200R and C4D164N at pH 8.0. Data were analyzed using the c(s) model in the software package SEDFIT. **D.** Native PAGE (7%) analysis of recombinant nonC4-NADP-ME variants. Top panel: Coomassie staining with 3 μg of C4- and nonC4-NADP-ME and 7.5 µg of mutant proteins per lane. Bottom panel: in-gel NADP-ME activity assay with 1 μg of C4 and nonC4 controls, and 2.5 µg of mutant proteins per lane. The SERVA native marker is shown on the left.

To further investigate the role of residues 163, 164, and 200 in NADP-ME oligomerization, we generated reverse mutants in nonC4-NADP-ME: nonC4R171T, nonC4N172D, and nonC4R208G. The nonC4R171T mutant, like the nonC4 isoform, exhibited an equilibrium between active tetramers and dimers, with a significantly higher proportion of dimers (Fig. 4D). In contrast, the nonC4N172D and nonC4R208G variants exhibited destabilized oligomeric structures, resulting in mixed states where only tetramers were enzymatically active (Fig. 4D). These variants also displayed a tendency to aggregate, which reduced the yield of soluble proteins. These findings highlight the critical roles of residues 163, 164, and 200 in C4-NADP-ME, as well as their corresponding ones in nonC4-NADP-ME, in regulating the formation and stabilization of NADP-ME oligomers.

Given the striking influence of G200 on the quaternary state of C4-NADP-ME, we further investigated its role in conjunction with N-terminal modifications. We produced two additional C4-NADP-ME variants: one with the N-terminal region of nonC4-NADP-ME extended by 15 residues and the G200R mutation (C4NnC4GR), and another with only the 15 residues at the N-terminus and the G200R mutation (C4N15GR) (Fig. 2E). In both cases, active dimers were observed (Fig. 2C), indicating that the G200R mutation alone is sufficient to drive the shift in oligomerization from tetramers to dimers in C4-NADP-ME. These results confirm that the N-terminal region of C4-NADP-ME is not directly involved in tetramerization.

### The N-terminal region is involved in the high turnover rate of C4-NADP-ME with specific adaptive substitutions linking quaternary state and malate affinity

In addition to exploring the implications for oligomerization state, we conducted a comprehensive analysis of the kinetic parameters for the resulting variants and mutants of both C4- and nonC4-NADP-ME isoforms. The kinetic analysis of the mutants indicated that elimination of the 15 residues at the N-terminus of nonC4-NADP-ME does not affect the kinetic parameters, similar to the nonC4DelN mutant (Table 1). Introducing the I148F mutation to the N-terminal deletion variant slightly increases the catalytic rate. Further mutations, including all 20 differentially substituted amino acids, further enhance the affinity for malate to values between those of the C4- and nonC4-NADP-ME (Table 1). These findings suggests that additional amino acids beyond those studied may contribute to the modulation of the C4-specific properties in maize C4-NADP-ME. Notably, while the 20 amino acids were identified as differentially conserved in both maize and sorghum nonC4 and C4 isoforms, further amino acid differences emerge when comparing only the maize nonC4 and C4 isoforms, potentially playing a role in the unique kinetic properties of maize C4-NADP-ME. It can also not be ruled out that the N-terminal region of the enzyme is involved in those properties.

**Table 1.**
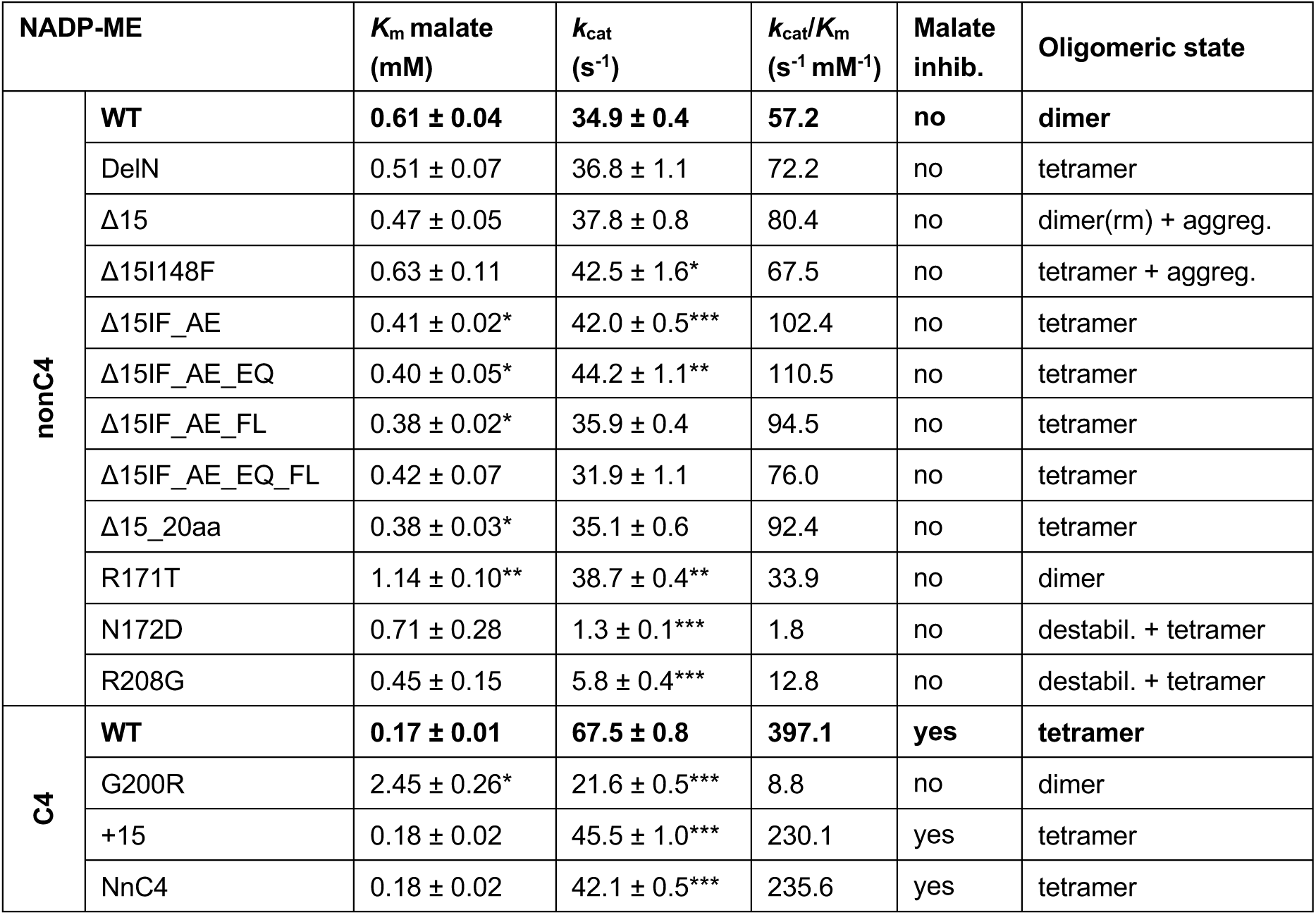
Kinetic parameters of recombinant C4- and nonC4-NADP-ME wild type and mutant variants. Data were adjusted to the Michaelis-Menten equation by non-linear regression with GraphPad Prism version 8.3.0 software (Boston, Massachusetts, USA). The parameters were determined at pH 8.0. The values represent mean ± standard deviation; n = three independent enzyme preparations, each measured in triplicate. For statistical analysis, a two-tailed t-test with Welch’s correction was performed, comparing nonC4- and C4-NADP-ME variants to the corresponding WT proteins. Asterisks indicate that the value is statistically significantly different from the corresponding WT. *P < 0.05; **P < 0.01; ***P < 0.001 (the P values are shown in Suppl. Table 2). aggreg. denotes aggregation. destabil. denotes destabilization of quaternary structure. Malate inhib. denotes inhibition by increasing concentrations of malate at pH 7.0. rm denotes slightly reduced mobility with respect to the nonC4-NADP-ME.

| NADP-ME | | $K_m$ malate (mM) | $k_{cat}$ ($s^{-1}$) | $k_{cat}/K_m$ ($s^{-1} \text{ mM}^{-1}$) | Malate inhib. | Oligomeric state |
| --- | --- | --- | --- | --- | --- | --- |
| nonC4 | WT | **0.61 ± 0.04** | **34.9 ± 0.4** | **57.2** | **no** | **dimer** |
| | DelN | 0.51 ± 0.07 | 36.8 ± 1.1 | 72.2 | no | tetramer |
| | Δ15 | 0.47 ± 0.05 | 37.8 ± 0.8 | 80.4 | no | dimer(rm) + aggreg. |
| | Δ15I148F | 0.63 ± 0.11 | 42.5 ± 1.6* | 67.5 | no | tetramer + aggreg. |
| | Δ15IF_AE | 0.41 ± 0.02* | 42.0 ± 0.5*** | 102.4 | no | tetramer |
| | Δ15IF_AE_EQ | 0.40 ± 0.05* | 44.2 ± 1.1** | 110.5 | no | tetramer |
| | Δ15IF_AE_FL | 0.38 ± 0.02* | 35.9 ± 0.4 | 94.5 | no | tetramer |
| | Δ15IF_AE_EQ_FL | 0.42 ± 0.07 | 31.9 ± 1.1 | 76.0 | no | tetramer |
| | Δ15_20aa | 0.38 ± 0.03* | 35.1 ± 0.6 | 92.4 | no | tetramer |
| | R171T | 1.14 ± 0.10** | 38.7 ± 0.4** | 33.9 | no | dimer |
| | N172D | 0.71 ± 0.28 | 1.3 ± 0.1*** | 1.8 | no | destabil. + tetramer |
| | R208G | 0.45 ± 0.15 | 5.8 ± 0.4*** | 12.8 | no | destabil. + tetramer |
| C4 | WT | **0.17 ± 0.01** | **67.5 ± 0.8** | **397.1** | **yes** | **tetramer** |
| | G200R | 2.45 ± 0.26* | 21.6 ± 0.5*** | 8.8 | no | dimer |
| | +15 | 0.18 ± 0.02 | 45.5 ± 1.0*** | 230.1 | yes | tetramer |
| | NnC4 | 0.18 ± 0.02 | 42.1 ± 0.5*** | 235.6 | yes | tetramer |

We thus analyzed the influence of the N-terminal region of C4-NADP-ME on its kinetic properties. Our findings reveal that the 15-residue insertion in the N-terminal region of C4-NADP-ME does not affect the enzymatic affinity for malate, as evidenced by the similar *K*_m_ values observed in the N-terminal modified variants, C4+15 and C4NnC4, compared to the wild-type enzyme (Table 1). However, these variants have a reduced turnover rate (*k*cat) compared to the wild type, suggesting that the N-terminal region is involved in modulating the catalytic properties of C4-NADP-ME.

Given that C4G200R exists as a stable dimer, we analyzed its kinetic parameters and compared them to those of the wild-type C4-NADP-ME (Table 1). At pH 8.0 C4G200R showed a 44-fold lower catalytic efficiency (*k*_cat_/*K*_m_ 8.8 s^−1^ mM^−1^) compared to wild-type C4-NADP-ME (*k*_cat_/*K*_m_ 397.1 s^−1^ mM^−1^). This substantial reduction is primarily due to a 3-fold decrease in turnover rate (C4G200R: *k*_cat_ = 21.6 ± 0.5 s^−1^; C4-NADP-ME WT: *k*_cat_ = 67.5 ± 0.8 s^−1^) and a 14-fold decrease in malate affinity (C4G200R: *K*_m_ = 2.45 ± 0.26 mM^−1^; C4-NADP-ME WT: *K*_m_ = 0.17 ± 0.01 mM^−1^). These results suggest that the G200 residue plays a crucial role in conferring the high malate affinity and catalytic rate characteristics of the C4 isoform, either directly or more likely through its contribution to tetramer formation. Interestingly, the C4G200R mutant, which forms a dimer, shows no inhibition by malate. In contrast, the C4-NADP-ME N-terminal mutants, C4+15 and C4NnC4, maintain a tetrameric structure and are inhibited by malate, similar to the wild-type isoform. These results provide the first evidence of a potential link between the tetrameric structure of C4-NADP-ME and its sensitivity to malate inhibition.

On the other hand, the tetrameric nonC4R208G variant exhibits a malate affinity comparable to the wild type but demonstrates extremely low catalytic efficiency (Table 1). This reduced enzymatic performance stems from the minimal fraction of active enzyme present in the tetrameric state (Fig. 4D). In a similar manner to the nonC4R208G variant, the nonC4N172D variant also displayed extremely low catalytic efficiency (Table 1), due to the minimal fraction of active enzyme existing as tetramers (Fig. 4D). However, this mutation did not significantly affect the affinity for malate. In contrast, the dimeric nonC4R171T variant exhibited a two-fold reduction in malate affinity while maintaining a turnover rate comparable to the wild-type enzyme, suggesting that R171 is likely involved in substrate affinity.

### Amino acid residues involved in nonC4- to C4-NADP-ME oligomeric state shifts localize at the dimer interface

To investigate how isoform-specific amino acid substitutions influence the oligomeric states of C4- and nonC4-NADP-ME, we generated dimeric models using AlphaFold3. The AlphaFold3 model for C4-NADP-ME demonstrated a remarkable structural alignment with the previously published crystal structure (PDB 5OU5; (Alvarez *et al*., 2019), validating our computational approach and establishing this crystal structure as the reference model for the C4 isoform (Fig. 5A).

**Figure 5.**
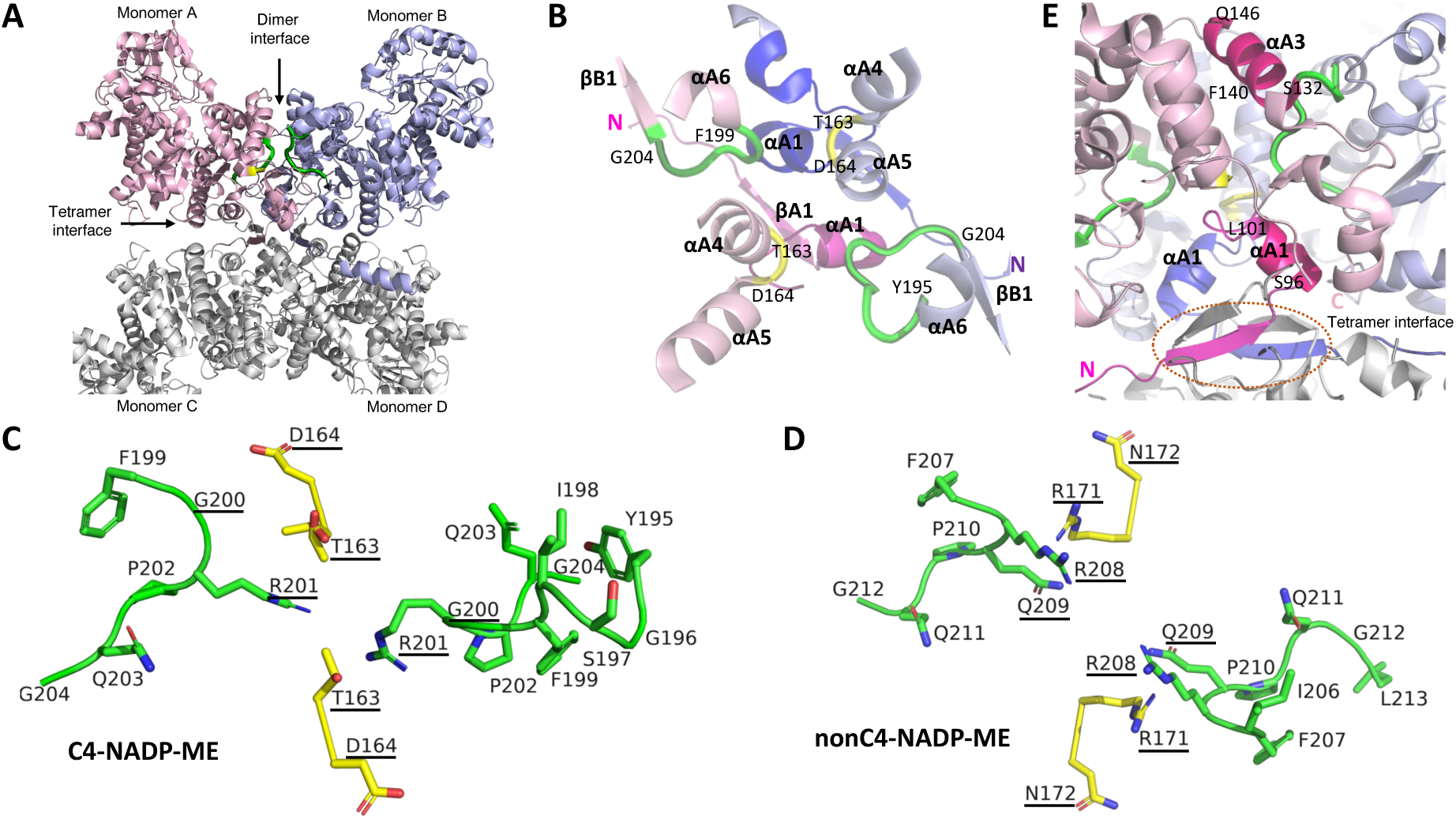
Critical loops involved in the differential oligomeric states of C4- and nonC4-NADP-ME isoforms. **A.** Side view of the C4-NADP-ME structure (PDB ID: 5OU5), showing monomer A in pink and monomer B in purple. The C-D dimer is depicted in grey. Loops containing residues T163 and N164 are highlighted in yellow, while loops containing residue G200 in green. **B.** Top view of the dimer interface in C4-NADP-ME. Loops formed by residues T163 and D164 are highlighted in yellow. Loops containing residues 199-FGRPQG-204 in monomer A and 195-YGSIFGRPQG-204 in monomer B are shown in green. N-termini are indicated. **C** and **D.** Structural comparison of the loops at the dimer interface in C4-NADP-ME (C) and nonC4-NADP-ME (D) The differentially conserved substitutions in the loops are underlined for each isoform. **E.** Detailed view of the C4-NADP-ME dimer interface, emphasizing the loops in proximity to helix αA3 and the N-terminal helix αA1. For simplicity, only residues 96-101 of αA1 and 132-146 of αA3 in monomer A are shown. N- and C-termini of monomer A are indicated. The N-terminal β-sheet structures, which mediate interactions among all four monomers at the tetrameric interface, are enclosed within an oval. In B and E secondary structures of monomer A are shown in pink and those of monomer B in purple.

The comparative structural analysis revealed that the dimer interface of both isoforms features four loops (Fig. 5A). Two of these loops, one in each monomer, contain residues T163 and D164 in C4-NADP-ME (highlighted in yellow in Fig. 5A and B) and the corresponding residues, R171 and N172, in nonC4-NADP-ME. In both isoforms, these loops connect helices αA4 and αA5 in each monomer (Fig. 5B). The remaining two loops, also present in each monomer, vary in sequence and length between the isoforms. In C4-NADP-ME, these loops contain residue G200 and bridge helix αA6 with the βB1 sheet (green loops, Fig 5B). Specifically, in monomer A, the loop comprises residues 199-FGRPQG-204, while in monomer B, it spans residues 195-YGSIFGRPQG-204 (Fig. 5B). Conversely, in nonC4-NADP-ME, the corresponding loops contain residue R208 and spans residues 207-FRQPQG-212 in monomer A and 206-IFRQPQGL-213 in monomer B.

Interestingly, the loops at the dimer interface of C4-isoform feature four differentially conserved amino acid substitutions (T163, D164, G200, and R201; Fig. 5C and D) and are in close proximity to helix αA3 (residues 132-146, Fig. 5E), which includes F140, the other differentially conserved amino acid substitution critical for stabilizing the tetrameric structure of C4-NADP-ME (Fig. 3A; (Alvarez *et al*., 2019)). In a previous work we have shown that F140 does not interact directly with the neighboring monomer, but the adjacent residue K138 and K139 form hydrogen bonds with E288 and G196 of the other monomer, respectively (Alvarez, 2019). Additionally, near the loops at the dimer interface lies the C-terminal loop (M627-R636) and the N-terminal helix αA1 (S96-L101) of each of the four monomers (Fig. 5E). Importantly, helix αA1 is positioned adjacent to the β-sheet structures that mediate interactions between all four monomers at the tetramer interface in C4-NADP-ME (Fig. 5E).

### Hot spot analysis identified residues 163 and 200 as key contributors to the stability of NADP-ME dimeric interfaces

Building on our previous findings, we employed the advanced machine learning tool KFC2 (Zhu and Mitchell, 2011) to identify critical interface residues, known as “hot spots” (HS), which are pivotal for binding affinity and specificity in protein-protein interactions. Using AlphaFold3, we constructed dimeric models of maize C4-NADP-ME, nonC4-NADP-ME, and the C4G200R mutant to analyze these HS.

A comparative analysis between nonC4- and C4-NADP-ME revealed 17 conserved HS between the isoforms (Suppl. Table 3). In addition, five residues (K147, Y155, N167, R171, and R208) were uniquely identified as HS in nonC4-NADP-ME, while two residues (R201 and C246) were exclusive HS in C4-NADP-ME (Suppl. Table 3). When comparing C4-NADP-ME with its C4G200R mutant, the G200R mutation introduced five new HS (K139, Y147, N159, R163, and R200) into the C4-NADP-ME structure. Remarkably, these newly identified HS correspond to the five HS uniquely found in nonC4-NADP-ME (Suppl. Table 3). Notably, N167, R171, and R208 in nonC4-NADP-ME are among the 20 differentially substituted amino acids between nonC4- and C4-NADP-ME. Of particular interest are R171 and R208, along with their corresponding residues T163 and G200 in C4-NADP-ME, which directly influence the oligomeric state of the proteins (Fig. 4B).

This analysis highlights the critical role of specific adaptive-substitutions in modulating monomer-monomer interactions and further demonstrates that the G200R mutation in C4-NADP-ME introduces key interface features resembling those of the nonC4 isoform.

### Amino acid interactions at the dimer interfaces of C4- and nonC4-NADP-ME

The presence of numerous differentially conserved amino acid substitutions at the dimer interface, which influence the oligomeric states of C4- and nonC4-NADP-ME, suggests that distinct interactions in this region may play a critical role in determining the different oligomeric conformations adopted by these isoforms. To investigate the interactions guiding the dimer conformation of monomers A and B in C4- and nonC4-NADP-ME, we conducted a detailed computational analysis using the PISA software (CCP4; (Krissinel and Henrick, 2007) and protein models generated by AlphaFold3. This analysis examined the residues involved in hydrogen bonds (Suppl. Table 4), salt bridges (Suppl. Table 5), and those contributing to the stabilization or destabilization of the dimer interface (Suppl. Table 6).

A comparison of interchain hydrogen bonds at the dimer interface reveals differences in the number and distribution of polar contacts between C4- and nonC4-NADP-ME. Specifically, C4-NADP-ME forms 17 polar contacts, while nonC4-NADP-ME forms 22. Of these, less than half are conserved between the isoforms (Suppl. Table 4). Different residues at the dimer interface contribute to the isoform-specific hydrogen bond networks: in C4-NADP-ME, the unique residues include Y98, L101, Q148, N159, G196, R201, and P202, while in nonC4-NADP-ME, the unique residues are R110, P136, N151, Y155, E170, S205, F207, R208, and R218 (Suppl. Table 4). These unique patterns of hydrogen bond interactions primarily involve residues located in helices at the dimer interface and the interconnecting loops (Fig. 5C and D). In C4-NADP-ME, Y98 and L101 are located in helix αA1, N159 in helix αA4, and R201 and P202 in the loop containing G200. In nonC4-NADP-ME, R110 in αA1 forms a hydrogen bond with E170 in αA4. Notably, the loop connecting αA4 and αA5 in nonC4-NADP-ME contains the residues R171 and N172, which correspond to T163 and D164 in C4-NADP-ME and contribute to the tetrameric structure organization (Fig. 4B). Furthermore, we observed a unique interaction in nonC4-NADP-ME that occurs within a key loop at the dimer interface, where R208 (which correspond to G200 in C4-NADP-ME) forms a hydrogen bond with Y155 (Suppl. Table 4). Finally, we observed that certain residues are conserved across the isoforms but form hydrogen bonds with different partners, such as Q152 and K139 in C4-NADP-ME and R131 in nonC4-NADP-ME (Suppl. Table 4).

The analysis of salt bridges revealed a high degree of conservation in the interactions at the dimer interfaces of both isoforms. The only observed differences are as follows: in C4-NADP-ME, there is a single salt bridge between D211 and R123, while in nonC4-NADP-ME, two salt bridges can be formed between the corresponding D219 and R131 residues (Suppl. Table 5). Notably, some residues involved in the salt bridges are located at the C-terminal end of the αA1 helix (R102 in C4-NADP-ME and R110 in nonC4-NADP-ME), while others reside within the αA3 helix (K138 in C4-NADP-ME and K146 in nonC4-NADP-ME). In particular, R110 in nonC4-NADP-ME forms a salt bridge with E170, which is adjacent to R171 and N172. These residues and the corresponding T163 and D164 in C4-NADP-ME are critical determinants in modulating the oligomeric states of the two isoforms (Fig. 4B).

Further investigation using PISA analysis (CCP4; (Krissinel and Henrick, 2007) into residues that either stabilize or destabilize the dimer interface revealed that P202 acts as a stabilizing contributor, while R201 serves as a destabilizing residue in C4-NADP-ME (Suppl. Table 6). Both residues are located within the G200 loop (Fig. 5B). In nonC4-NADP-ME, P136 was identified as a stabilizing residue, whereas R171 was found to be destabilizing.

In summary, this analysis underscores the importance of specific dimer interface residues, particularly within the key loops, in modulating the oligomeric states of C4- and nonC4-NADP-ME isoforms. The G200 loop (with residues G200, R201 and P202) in C4-NADP-ME and the loop containing R171 in nonC4-NADP-ME appear to play pivotal roles in these processes.

### Crystallographic structure of maize C4G200R mutant and oligomeric state analysis by cryo-EM

Due to the critical role of G200 in shifting the oligomerization state of C4-NADP-ME, we solved the crystal structure of C4G200R at 2.7 Å resolution at pH 4.8 (see Suppl. Table 7 for data collection and refinement statistics). The structure features good stereochemistry, with the asymmetric unit containing surprisingly a single homotetrameric molecule, best described as a dimer of dimers. The polypeptide chains exhibit continuous electron density consistent with the resolution of the diffraction data, except for the first 11-25 residues (depending on the monomer analyzed), which could not be resolved, likely due to increased mobility within the crystal.

The C4G200R structure closely resembles wild-type C4-NADP-ME in both architecture and organization (Fig. 6A). Superimposition of the tetramers yields a root-mean-square deviation (RMSD) of only 1.15 Å across 2,128 aligned Cα atoms, underscoring their structural similarity. Additionally, dimers A-B and C-D are tilted relative to each other in a manner similar to the wild-type enzyme (Fig. 6B). The four C4G200R monomers exhibit highly similar atomic positions, with RMSD values ranging from 0.21 to 0.42 Å for over 550 aligned Cα atoms. The buried surface area for the tetramer is 19,360 Å², while the A-B and C-D dimers have buried areas of 6,130 and 5,940 Å², respectively. The model includes pyruvate molecules at each of the four active sites (Fig. 6A). No evidence of bound NADP was observed in the density.

**Figure 6.**
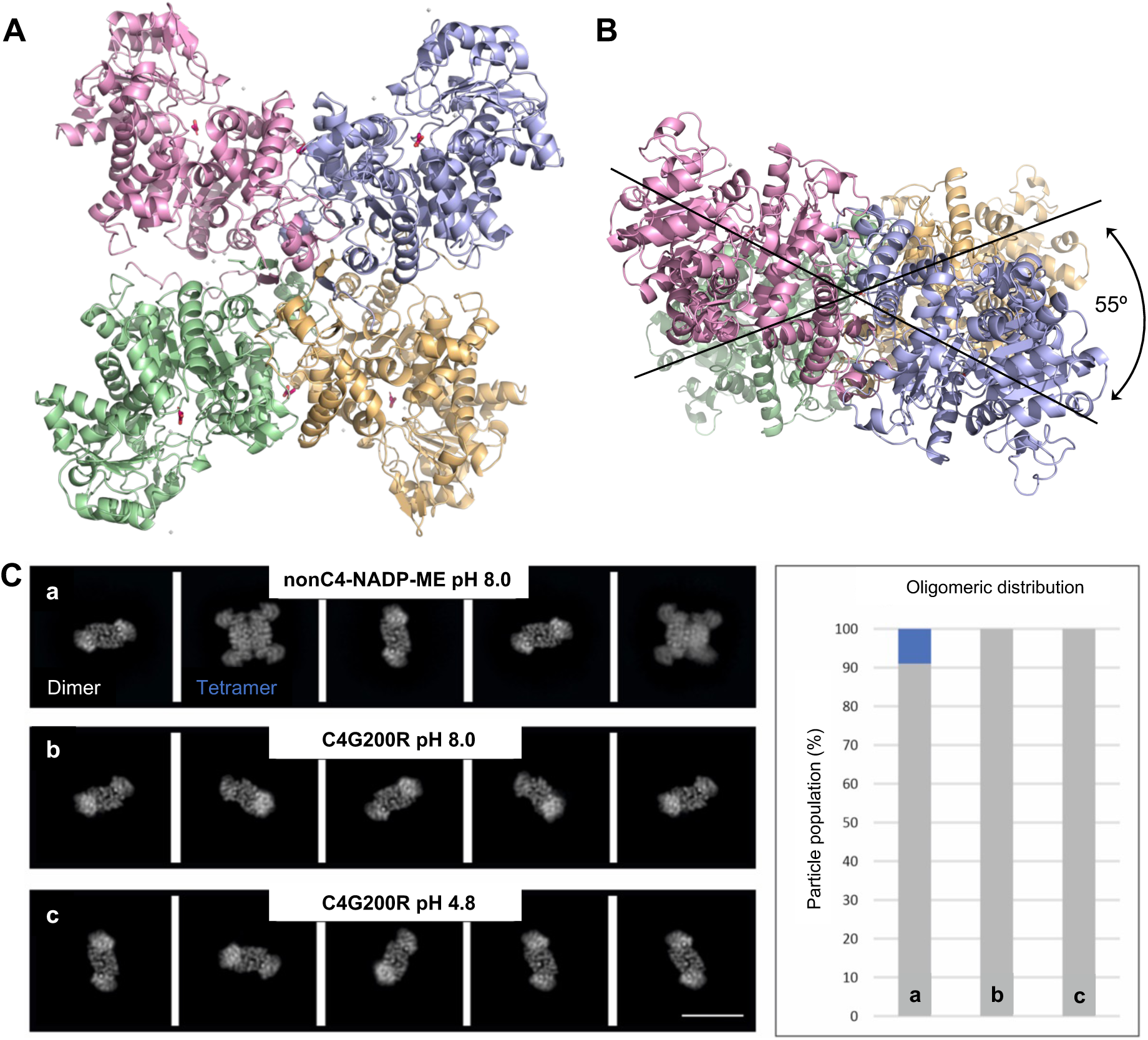
Crystallography and cryo-EM analysis. **A.** Cartoon illustration of the crystallographic structure of C4G200R. Monomer A (pink) and monomer B (purple) represent the dimer; likewise, monomer C (green) and monomer D (yellow). Pyruvate molecules at the four active sites and two dimeric interfaces are displayed as stick models in red. **B.** Top view of C4200R, displaying the tilted arrangement of the dimers relative to each other. **C.** 2D cryo-EM analysis of nonC4-NADP-ME and C4G200R. Representative 2D class averages of nonC4-NADP-ME at pH 8.0 (a), C4G200R at pH 8.0 (b) and C4-NADP-ME-G200R at pH 4.8 (c), respectively (Scale bar: 110 Å). The distribution (%) of oligomeric state (dimer; gray, tetramer; blue) of each sample is shown based on the number of single particles showing high resolution features for the respective oligomeric state.

X-ray crystallography requires proteins to form a highly ordered crystal lattice, where proteins are tightly packed. This lattice can sometimes impose or stabilize a specific oligomeric state that may not be the predominant form in solution (Sala *et al*., 2020). As a result, crystallization can promote interactions that favor higher order oligomers - such as the tetramers observed - that may not occur naturally or may be less stable in solution, which may explain the discrepancy with the native PAGE and AUC results. To better capture the protein’s oligomeric state in a more physiological setting, we also conducted cryo-electron microscopy (cryo-EM), which visualizes proteins in a near-native, frozen-hydrated state. 2D single particle cryo-EM analysis of C4G200R (pH 4.8 and 8.0) and nonC4-NADP-ME (pH 8.0) revealed that C4G200R forms exclusively dimers under both pH conditions, whereas nonC4-NADP-ME additionally contains a small particle population (10%) of tetramers, which is consistent with the native PAGE results (Fig. 6C).

The observation of C4G200R forming a tetramer in X-ray crystallography, but a dimer in cryo-EM under similar conditions, as well as in native PAGE, suggests that the tetrameric assembly is likely an artifact induced by crystal packing forces rather than a physiologically relevant oligomeric state. Furthermore, the ability of the protein to adopt different oligomeric states indicates that the quaternary structure is sensitive to experimental conditions, including protein concentration, ionic strength, and buffer composition.

### Differential spatial positioning of residues at loop containing G200 in C4G200R promotes its dimerization

The G200R mutation is located within the loop connecting domain A (αA6) and domain B (βB1) (Fig. 7A). A comparison of the crystallographic structures of C4G200R and C4-NADP-ME reveals significant differences at the main chain positions of residues 199 and 200 (Fig. 7A). In the mutant structure, the altered positioning of these residues affects interactions at the dimer interface, as identified through PISA analysis (Suppl. Table 8). Specifically, the hydrogen bond interaction between Y98 (near the N-terminus of one monomer) and P202 (part of the “G200 loop” and adjacent to R201, a key destabilizing residue in C4-NADP-ME) is disrupted in the G200R mutant (Fig. 7B and Suppl. Table 8).

**Figure 7.**
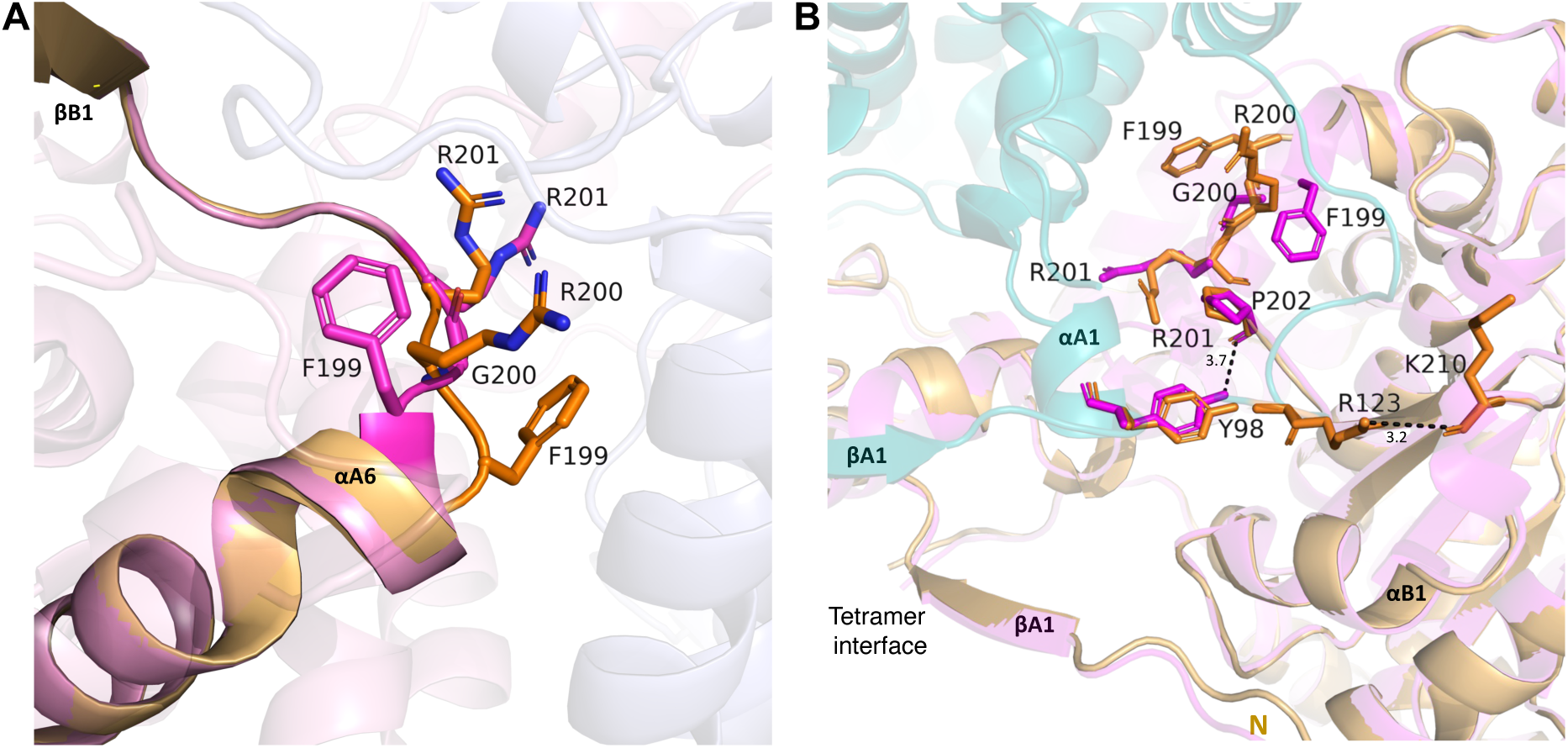
Key structural differences between C4-NADP-ME and C4G200R. **A.** Comparative structural representation showing significant changes in the 199-FGRPQG-204 loop between maize C4-NADP-ME and C4G200R. Residues F199, R200 and R201 in C4G200R are depicted as orange sticks, while F199, G200 and R201 in C4-NADP-ME are represented as pink sticks. **B.** Relative positioning of the differential interactions identified by PISA analysis (Suppl. Table 8) between C4-NADP-ME and the C4G200R.

In addition, two new hydrogen bonds form in the G200R mutant between R123 and K210 (Fig. 7B), reflecting structural changes resulting from the G200 mutation. This, together with the HS analysis showing that the incorporation of R200 in C4-NADP-ME introduces HS residues unique to nonC4-NADP-ME (Suppl. Table 3), highlights the pivotal role of the G200-containing loop and its specific interactions with the N-terminal region in mediating isoform-specific interactions and regulating the oligomerization state of the plastidic NADP-ME isoforms.

### The C-terminal ends of C4- and nonC4-NADP-ME play a key role in the oligomeric state stability

Structural analyses of C4- and nonC4-NADP-ME isoforms revealed that the loops containing residues G200, T163, and D164 are located near the N- and C-terminal regions of the monomers, specifically at the tetrameric interface (Fig. 5C, Fig. 8A). To explore potential key interactions in these regions, we conducted a detailed search for hydrogen bonds using the PyMOL structural modeling program.

**Figure 8.**
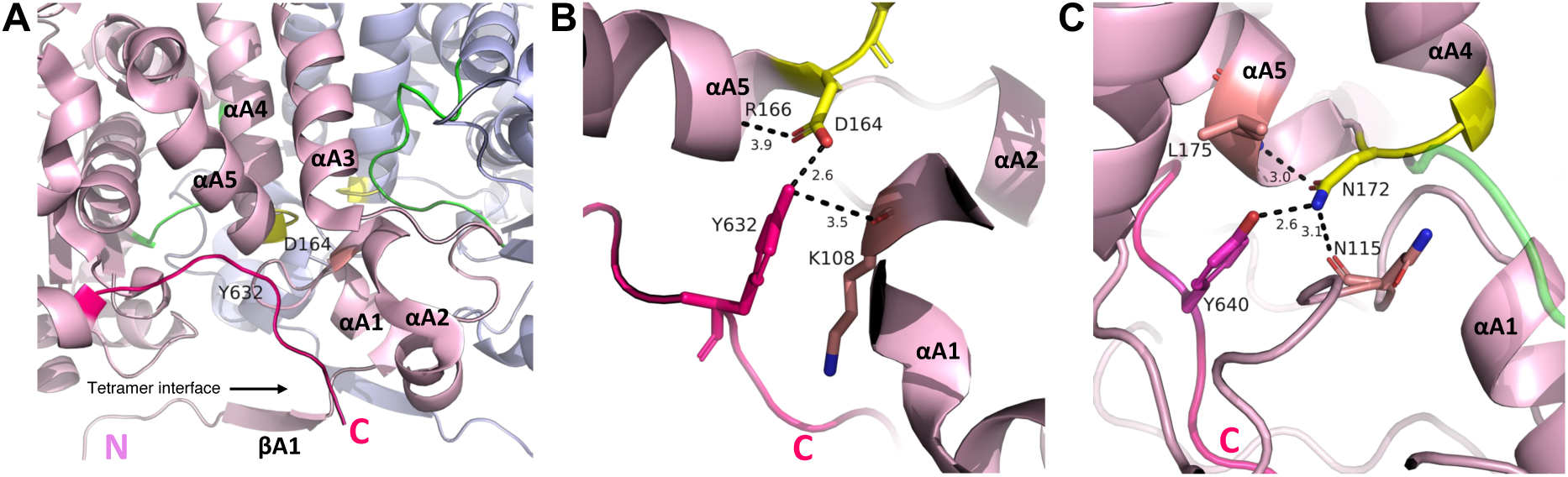
Interactions of the C-terminal end of C4- and nonC4-NADP-ME. **A.** Relative positioning of the N- and C-terminal ends, highlighting key loops containing residues T163 and D164 (yellow) and G200 (green) in monomers A and B of C4-NADP-ME. For simplicity, only monomers A (pink) and B (purple) are shown. The N- and C-termini, along with secondary structures near the tetramer interface, are indicated for monomer A. **B.** Polar interactions of D164 with R166, Y632, and K108 at the tetrameric interface of C4-NADP-ME. **C.** Polar interactions of N172 with L175, Y640, and N115 at the tetrameric interface of nonC4-NADP-ME. In panels B and C, black dashed lines indicate polar bonds between the interacting functional groups, with bond distances denoted in Å.

In the C4-NADP-ME isoform, a critical network of polar interactions was identified. The carboxylic side chain of residue D164 simultaneously forms two hydrogen bonds: one with the backbone of residue R166 and another with the hydroxyl group of Y632. Additionally, Y632 establishes a hydrogen bond with the main chain carbonyl group of K108, located in the loop connecting αA1 and αA2 at the N-terminal end (Fig. 8B). In contrast, the nonC4-NADP-ME isoform exhibits other interactions due to the presence of residue N172 as positionally homolog of D164 (C4-adaptive substitution) in the C4 isoform. The carbonyl group of the N172 side chain interacts with the amine backbone of L175 (which corresponds to L167 in C4-NADP-ME), while its amine group forms simultaneous hydrogen bonds with Y640 (which corresponds to Y632 in C4-NADP-ME) and the backbone carbonyl group of N115 (which corresponds to N107 in C4-NADP-ME) (Fig. 8C).

These distinct interactions in both isoforms bring the N- and C-termini of each monomer closer together, clustering them centrally at the tetrameric interface (Fig. 8A). Notably, we showed that in C4-NADP-ME, the D164N mutation altered the protein oligomeric state, whereas the reciprocal N172D mutation in nonC4-NADP-ME disrupted oligomeric stability, leading to a heterogeneous population of oligomeric states with enzymatic activity preserved exclusively in the tetrameric configuration (Fig. 4B and D). These results indicate that the ionic interactions of these residues are critical for maintaining the oligomeric structure.

To further investigate the functional role of the C-terminal end in protein stability, we generated the C4Y632F mutant by replacing tyrosine with phenylalanine, thereby removing the hydroxyl group essential for interactions with D164. Biochemical analysis revealed that this mutation disrupted the stability of the tetrameric assembly, leading to the formation of a mixture of oligomeric states (Fig. 4D). Enzymatic activity was retained solely in the tetrameric configuration. Notably, this instability closely resembled the effects of the F140I mutation (Fig. 3A).

These findings highlight the pivotal role of C-terminal interactions in preserving the structural integrity of both C4- and nonC4-NADP-ME isoforms. The interactions between C-terminal residues and key C4-adapted residues at the dimer interface seems to be crucial for the differential oligomeric stability and enzymatic functionality of these isoforms.

## Discussion

A diverse array of mechanisms defines the oligomerization state of a protein, including domain swapping, ligand-induced dimerization, point mutations at dimer interfaces, post-translational modifications, and small amino acid insertions or deletions (Sun *et al*., 2002, Matsumiya *et al*., 2003, Hashimoto and Panchenko, 2010). In this study, biochemical and structural analyses of recombinant NADP-ME isoforms and their mutants, combined with predictive algorithms, revealed that specific amino acid substitutions at the dimer interface and distinct N-terminal features selectively influence the oligomeric structures of maize plastidic C4-photosynthetic and housekeeping NADP-ME isoforms. Our findings support an evolutionary model for the divergent evolution of oligomeric structures in plastidic NADP-ME isoforms in maize and sorghum.

### Stabilization of the C4-NADP-ME tetrameric form was achieved by mutations in the dimer interphase

The evolutionary transition to C4 photosynthesis has been accompanied by massive upregulation of genes encoding C4 pathway enzymes, including NADP-ME (Aubry *et al*., 2011, Maier *et al*., 2011). Since highly expressed proteins face increased risks of aggregation and misfolding due to their abundant presence in the cellular environment, they typically evolve specific sequence and structural features that enhance their stability (Tartaglia *et al*., 2007). The evolutionary transition to a C4-optimized NADP-ME occurred under selective pressure (Christin *et al*., 2009). We propose that certain adaptive amino acid changes promote a particularly stable tetrameric assembly at physiological pH, as oligomerization reduces the solvent-exposed surface area per monomer, enhancing stability and minimizing the risk of protein aggregation despite its high cellular concentration (Miller *et al*., 1987, Jones and Thornton, 1995). This optimization of the quaternary structure might also have favored the emergence of unique biochemical properties to regulate carbon flux in the C4 fixation pathway. These adaptations include increased enzyme turnover rates and enhanced malate affinity, along with pH-sensitive malate inhibition occurring between pH 7.0-7.5. The functional importance of the C4-NADP-ME quaternary structure is highlighted by the dimeric C4G200R mutant, which loses the characteristic malate inhibition (Table 1).

Consistent with this, we identified strictly differentially substituted amino acids along the dimer interface that, when mutated to those found in the housekeeping nonC4 isoform, destabilize the quaternary structure of C4-NADP-ME, converting it into a dimer (G200R), shifting it to a dimer-tetramer equilibrium (T163R, D164N), or resulting in a mixture of oligomeric states from tetramer to monomer (F140I; (Alvarez *et al*., 2019)). The same residues also affect the quaternary structure of nonC4-NADP-ME when mutated to those found in the C4 isoform. All this indicates that the adaptive substitutions acquired by C4-NADP-ME stabilize the dimer interface, enhancing dimer-dimer interactions and thereby supporting a stable tetrameric structure. Furthermore, specific interactions involving the strictly conserved residues D164 in C4-NADP-ME and its positional homolog N172 in nonC4-NADP-ME, as well as the C-terminal residue Y632 in C4-NADP-ME and its positional homolog Y640 in nonC4-NADP-ME, indicates that the C-termini of both isoforms play a critical role in maintaining the oligomeric stability of these isoforms. This mechanism is analogous to the human mitochondrial NAD(P)-ME, where the oligomeric state is stabilized by a C-terminal extension (Xu *et al*., 1999).

### The N-terminus restricts tetramer formation in nonC4-NADP-ME

Small sequence insertions or deletions can serve as an important evolutionary mechanism affecting oligomerization, potentially altering stability of oligomer and enabling new functional specificities (Hashimoto and Panchenko, 2010). For instance, insertions/deletions can introduce/eliminate secondary structural elements that either facilitate or hinder protein-protein interactions (Jiang and Blouin, 2007, Akiva *et al*., 2008). In maize and sorghum, the dimeric nonC4-NADP-ME has a 15-residue longer N-terminal region compared to the tetrameric C4 isoform. This extended N-terminal sequence, partially inherited from the ancient cytosolic isoform and partially acquired through the addition of the transit peptide, was later eliminated in the evolution of the C4 isoform.

We found that removing only these 15 residues from the N-terminus of nonC4-NADP-ME (nonC4Δ15) altered its mobility in native PAGE and promoted aggregation, indicating that this region is crucial for maintaining quaternary structure stability. Truncation of the N-terminus until the midpoint of the 15 residues (nonC4DelN) induced tetramerization, suggesting the presence of an inhibitory element within this region that prevents specific dimer-dimer interactions in the nonC4-NADP-ME. Consistently, adding the nonC4-NADP-ME N-terminal extension to the maize cytosolic isoform Cyt2 induced its dimerization, confirming that the residues AAGVV within the extended N-terminus of nonC4-NADP-ME disrupt dimer-dimer interactions. Hashimoto and Panchenko (2010) reported that 59% of disrupting regions in proteins form loops. In nonC4-NADP-ME, the exact structural basis for tetramer prevention remains uncertain, as this N-terminal region is not resolved in any crystallographic structure. Interestingly, the N-terminal region of pigeon liver NADP-ME has also been shown to affect oligomerization (Chou *et al*., 1997). In Choús study, the wild-type enzyme predominantly formed tetramers on native PAGE, while removal of the first three residues shifted the oligomeric equilibrium toward dimers, and further deletion of the first 16 residues resulted exclusively in dimers.

Our analysis revealed that modifying the N-terminal region of C4-NADP-ME by adding the 15 extra residues from nonC4-NADP-ME, or even replacing the entire N-terminal sequence with that of nonC4-NADP-ME, did not affect its oligomeric state. This suggests that the N-terminal region does not play a role in oligomerization in the C4 isoform. When we introduced mutations into nonC4Δ15, specifically targeting amino acids that are differentially substituted between maize and sorghum C4- and nonC4-NADP-ME (Alvarez *et al*., 2019), we observed stabilization of the tetrameric structure. In contrast, the corresponding positional mutations in nonC4-NADP-ME destabilize its dimeric form, resulting in a mixture of oligomeric states. Significantly, our kinetic analyses demonstrate that the N-terminal 15-residue deletion in C4-NADP-ME plays a crucial role in optimizing catalytic efficiency. This is evidenced by the reduced turnover rate values observed in the C4+15 and C4NnC4 variants compared to the wild type enzyme (Table 1). These findings reveal a dual evolutionary adaptation: while specific amino acid substitutions evolved to maintain tetrameric stability following the ancestral N-terminal deletion, this same deletion proved instrumental in achieving the enhanced catalytic rates characteristic of C4 metabolism.

### Which factors may have driven the divergent evolution of oligomeric structures in plastidic NADP-ME isoforms in maize and sorghum?

In maize and sorghum, nonC4-NADP-ME is expressed across different cell types and organs (Fig. 1B; (Maurino *et al*., 2001, Alvarez *et al*., 2013), while C4-NADP-ME is exclusively localized to BSC chloroplasts (Maurino *et al*., 1997, Swift *et al*., 2024). In C4 plants, unique gene or transcript sequences called *cis*-elements interact with *trans*-acting factors to control the accumulation of proteins essential for C4 photosynthesis within BSCs (Patel *et al*., 2004, Patel *et al*., 2006, Brown *et al*., 2011). Similar regulatory elements, located in the 5′ region of coding sequences, have been adapted across angiosperm monocots and eudicots to recognize *trans*-acting factors only found in C4 plants that confer BSC specificity (Brown *et al*., 2011, Kajala *et al*., 2012, Swift *et al*., 2024). This exemplifies the parallel evolution of gene regulation of C4-NADP-ME in maize and C4-NAD-ME in Cleome, where identical *cis*-elements at conserved positions were recruited for BSC-specific expression (Brown *et al*., 2011). Interestingly, these *cis*-elements are also present in C3 plant orthologs, where the necessary *trans*-factors are absent, suggesting that the *trans*-factors have been repeatedly recruited throughout C4 evolution to utilize pre-existing *cis*-elements in ancestral C4 genes (Brown *et al*., 2011, Kajala *et al*., 2012, Williams *et al*., 2012).

Given that both C4- and nonC4-NADP-ME evolved from ancient plastidic precursor in maize and sorghum, and this ancient plastidic precursor from an ancient cytosolic isoform related to the Cyt2 gene in maize, we analyzed whether nonC4-NADP-ME and Cyt2 contain the conserved *cis*-regulatory elements identified by Brown *et al*. (2011). By aligning C4 and nonC4-NADP-ME from maize and sorghum, as well as Cyt2 from maize and C3 and C4-NAD-ME sequences, we found that nonC4-NADP-ME in both species and maize Cyt2 preserve all *cis*-elements at conserved positions, suggesting that these elements were inherited from a common cytosolic ancestral gene (Suppl. Fig. 4). However, these elements are located farther upstream from the ATG start codon in nonC4-NADP-ME, due to the additional 45 nucleotides in the 5′ region and very near to the ATG in Cyt2 (Suppl. Fig. 4). Brown *et al*. (2011) shown that the correct positioning of these *cis*-elements is important for the BSC-specific *NAD(P)-ME* gene expression. Transcripts encoding nonC4-NADP-ME were found in various maize tissues, yet were absent in BSCs (Tausta *et al*., 2002). The increased distance between the *cis*-regulatory elements and the ATG start codon may therefore reduce or even suppress gene expression in these cells. However, it is also possible that additional factors, likely located in the promoter region, contribute to the broader expression pattern of nonC4-NADP-ME. In support of this, recent research demonstrated that sorghum DOF (DNA-binding with one finger) transcription factors can activate BSC expression by binding to the sorghum C4-NADP-ME promoter, which in maize and sorghum is enriched in DOF motifs (Borba *et al*., 2023, Swift *et al*., 2024). This suggests that a combination of regulatory elements may play a role in modulating the expression of the plastidic NADP-ME isoforms in different cell types.

Beyond potentially disrupting BSC-specific expression, we demonstrated that the first few nucleotides encoding the residues AAGVV in the extended N-terminal region of nonC4-NADP-ME enable this isoform to form a dimeric structure. This dimerization may have allowed the isoform to acquire new functional roles within the unique biochemical environment of C4 plants. While the exact role of plastidic nonC4-NADP-ME remains unclear, our findings suggest a functional specialization distinct from that of its C3 plant orthologs. Further research is needed to fully elucidate the biological function of nonC4-NADP-ME in C4 plants.

### Model for the divergent evolution of oligomeric structures in plastidic NADP-ME isoforms in maize and sorghum

Our findings support the following model for the evolution of distinct oligomerization structures in plastidic NADP-ME isoforms in plants of the C4 Andropogoneae lineage (Fig. 9). In the C3 ancestor, a gene duplication event involving an ancient cytosolic isoform (ancient Cyt2) gave rise to an ancient plastid-localized form, facilitated by the acquisition of an N-terminal transit peptide. This N-terminal region contained amino acids critical for dimerization. Moreover, the ancient plastidic isoform also contained *cis*-elements required for expression in BSCs inherited from the ancient cytosolic isoform, although these elements may have been inactive in the C3 context without the appropriate *trans*-acting factors (Brown *et al*., 2011).

**Figure 9.**
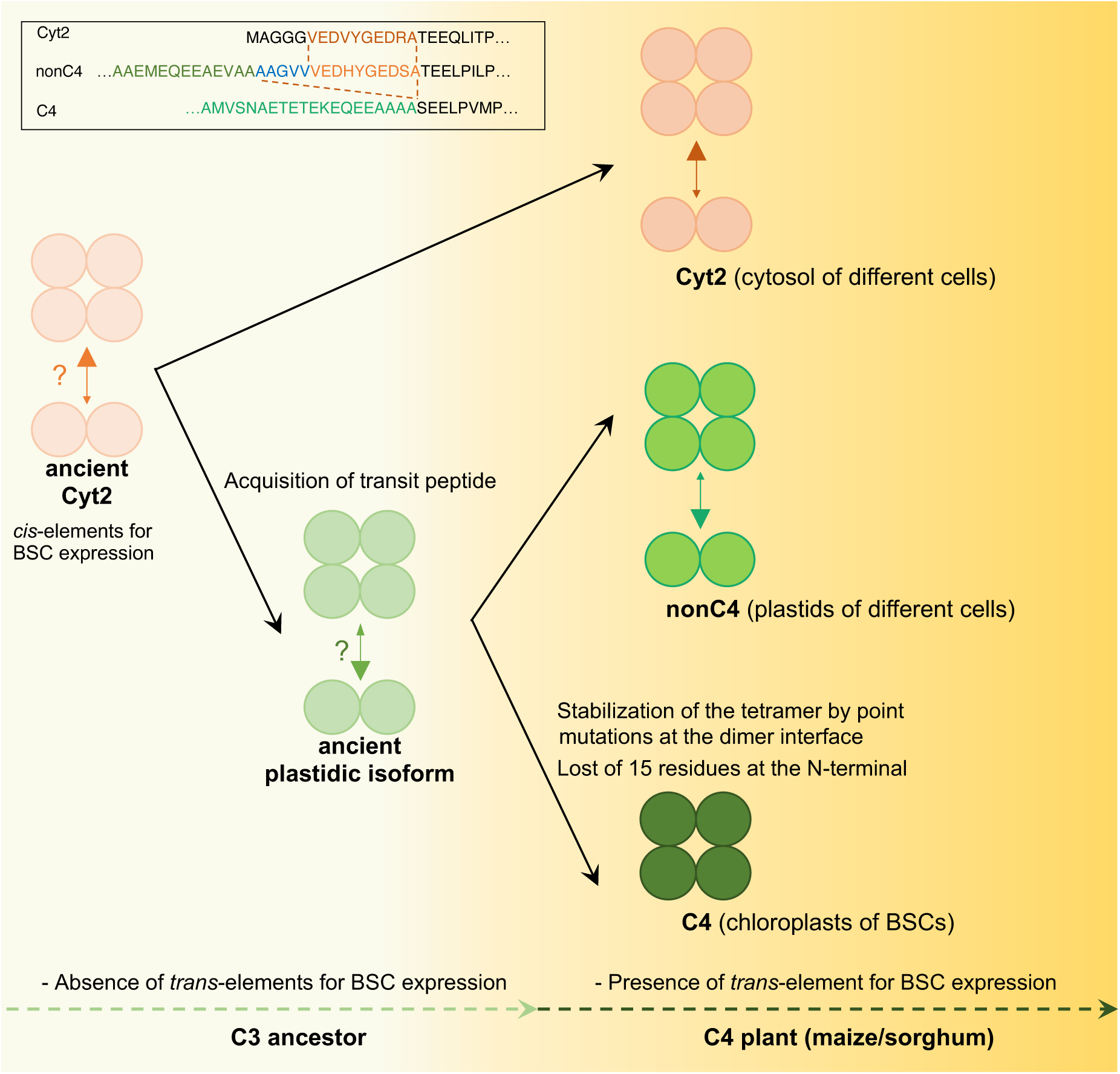
Proposed evolution of the oligomeric assembly of plastidic housekeeping (nonC4) and photosynthetic (C4) NADP-ME in maize and sorghum. The inset in the upper left corner shows the N-terminal sequences of maize Cyt2, nonC4-NADP-ME and C4-NADP-ME isoforms. Arrow sizes correspond to the relative prevalence of each oligomeric state, with question marks indicating the most probable state of the ancient isoforms. The scheme was designed as a free-form artistic illustration, with the lines not representing branch lengths.

With the evolution of C4 photosynthesis, a subsequent duplication of the ancestral plastidic isoform gave rise to both the C4- and nonC4-NADP-ME isoforms, each retaining the transit peptide and *cis*-elements for BSCs expression. The C4-NADP-ME, with a central role in the C4 photosynthetic pathway, adapted by evolving high catalytic efficiency, enhanced affinity for malate, inhibition by malate at pH 7.0-7.5, and a stable tetrameric structure. These features were driven by specific amino acid substitutions and likely to changes at the N-terminal region of C4-NADP-ME. It remains to be determined whether the 15-residue deletion in the N-terminal region of C4-NADP-ME is related to the positioning of the *cis*-regulatory elements, which are essential for its preferential expression in BSCs.

### C4-NADP-ME adaptations: lineage-specific rather than a universal solution

In the early 1990s, Iglesias and Andreo (Iglesias and Andreo, 1990) proposed a regulatory mechanism for C4 pathway flux based on their observations using C4-NADP-ME isolated from sugarcane leaves. Theirs *in vitro* studies showed that the oligomerization state of the enzyme varied with pH, leading to the hypothesis that such pH-dependent structural changes might modulate enzyme activity, and in turn carbon flux between day and night in C4 plants. While this mechanism may hold true for sugarcane, our experimental findings with maize and sorghum C4-NADP-ME suggest that it is not universally applicable. Under physiologically relevant conditions, the maize and sorghum enzymes maintain a remarkably stable tetrameric structure, showing no evidence of pH-dependent oligomeric transitions. This species-specific difference aligns with our previous analysis, which revealed numerous lineage-specific amino acid substitutions in C4-NADP-ME across different C4 lineages (Alvarez *et al*., 2019). These findings suggest that C4-NADP-ME optimization has followed distinct evolutionary trajectories in different plant lineages, likely adapting to species-specific physiological contexts (Eckardt *et al*., 2024). Such divergent evolution carries significant implications for metabolic engineering, particularly in efforts to engineer C4 photosynthesis into C3 plants (Ermakova *et al*., 2021). Successfully implementing this process may necessitate precise matching of C4-NADP-ME variants to their target cellular environment, as enzymes optimized for one species might exhibit suboptimal performance in another.

## Methods

### Generation of expression constructs of C4-NADP-ME and nonC4-NADP-ME variants

To generate recombinant maize wild-type C4- and nonC4-NADP-ME, we used the expression vectors *pET16b::ZmC4-NADP-ME* and *pET28::ZmnonC4-NADP-ME*, which were constructed in our previous work (Alvarez *et al*., 2019). Expression vectors for various mutant variants of C4- and nonC4-NADP-ME were produced either through commercial gene synthesis by BioCat (Heidelberg, Germany) or by site-directed mutagenesis (Suppl. Table 9). In the case of commercial gene synthesis, the cDNAs were cloned into pET16b via NdeI/XhoI restriction sites.

### Site directed mutagenesis

Some mutant variants were generated by introducing point mutations into the expression vectors *pET16b::ZmC4-NADP-ME*, *pET28::ZmnonC4-NADP-ME*, or other vectors synthesized commercially (see Suppl. Table 9 for detailed information). To accomplish this, the entire expression construct was amplified by PCR using Phusion High-Fidelity DNA Polymerase (Thermo Fisher Scientific, Waltham, USA) and overlapping primers containing the desired nucleotide changes (see Suppl. Table 10 for a list of primers used for site-directed mutagenesis). After PCR amplification, the template plasmid was removed by digestion with DpnI (Thermo Fisher Scientific, Waltham, USA) at 37 °C for 1 hour. The digested PCR product was then purified using the Monarch PCR & DNA Cleanup Kit (New England Biolabs, Ipswich, USA) and used to transform *E. coli* DH5α. After plasmid purification, successful site-directed mutagenesis was confirmed by sequencing the expression vectors through EZ-seq services (Macrogen Europe, Amsterdam, The Netherlands).

### Generation of expression constructs of Cyt2 variants via Gibson assembly

To generate recombinant maize Cyt2 (B6TVG1), we used the *pET32a::ZmCyt2-NADP-ME* vector (Alvarez *et al*., 2013) as a template for PCR-based fragment construction, following the method described by Gibson *et al*. (2011). For protein expression, Cyt2 and its N-terminal variants were cloned into the pET16b expression vector. For the Cyt2 construct, Gibson overhangs were incorporated in a single PCR reaction using primers complementary to both the fragment and the destination vector (see Suppl. Table 10 for a list of primers). The resulting PCR product was purified using the Monarch PCR & DNA Cleanup Kit (New England Biolabs, Ipswich, USA). For the Cyt2N variant, the first 14 residues of nonC4-NADP-ME were fused to the N-terminal of Cyt2. This additional sequence was introduced in two consecutive PCR reactions with Phusion High-Fidelity DNA Polymerase (Thermo Fisher Scientific, Waltham, USA). After each PCR, the product was purified using the Monarch PCR & DNA Cleanup Kit. Primers were designed with one portion annealing to the PCR fragment and the other containing an overhang corresponding either to the additional nucleotide sequence or to the destination vector (see Suppl. Table 10 for a list of primers). In the first PCR, part of the additional sequence was incorporated, while the second PCR introduced the remaining sequence along with the Gibson overhangs.

In the Cyt2DelN variant, the first 19 residues of nonC4-NADP-ME were fused to and partially replaced the N-terminal region of Cyt2. This nucleotide modification was introduced through three consecutive PCR reactions using Phusion High-Fidelity DNA Polymerase (Thermo Fisher Scientific, Waltham, USA). After each PCR step, the resulting fragments were purified using the Monarch PCR & DNA Cleanup Kit. In the first PCR, the original N-terminal sequence was mutated by introducing two nucleotide substitutions and adding three additional nucleotides. In the second PCR, as described for the Cyt2N variant, the first portion of the additional nucleotide sequence was added. The purified fragment was then inserted into a TOPO vector using the Zero Blunt TOPO PCR Cloning Kit (Thermo Fisher Scientific, Waltham, USA). After amplification in *E. coli* DH5α, the vector was sequenced to confirm the successful mutation of the N-terminus. The TOPO vector was then used as a template for the final PCR step, where the remaining nucleotide sequence and Gibson overhangs were added.

The empty pET16b vector was linearized with XhoI, and the purified fragments were inserted into the vector using Gibson isothermal assembly (Gibson, 2011). The assembly reaction was carried out using reagents from Thermo Fisher Scientific (Waltham, USA) and New England Biolabs (Ipswich, USA). The resulting construct was transformed into *E. coli* DH5α, and correct insertion into the pET16b vector was verified by sequencing.

### Heterologous expression of NADP-ME variants

Expression constructs were transformed into chemically competent *E. coli* Rosetta2 (DE3; pLysSRARE2) cells (Merck, Darmstadt, Germany) or ArcticExpress (DE3) cells (Agilent Technologies, Santa Clara, USA). Transformed cells were plated on lysogeny broth (LB) agar plates containing 100 µg/ml ampicillin (for pET16b) or 50 µg/ml kanamycin (for pET28b) to select for the presence of the respective plasmids.

For the expression of His-tagged proteins, pre-cultures were grown overnight in LB medium containing the appropriate antibiotics at 37°C shaking at 140 rpm. The following day pre-cultures were used to inoculate 400 ml of main cultures and further grown under the same conditions until an absorbance at λ = 600 nm (OD600) of 0.4-0.6 was reached, at which point protein expression was induced by adding 1 mM isopropyl-ß-D-thiogalactopyranoside (IPTG).

Cells were harvested 2 hours after induction. For Arctic Express cells, cultures were grown at 30 °C until an OD600 of 0.4-0.6 was reached, then cooled to 11 °C, and protein expression was induced with 1 mM IPTG. Cells were cultured at 11 °C for 24 hours before harvesting. To harvest the cells, cultures were centrifuged at 6000 × g for 10 minutes at 4°C, and the pellets were stored at −20 °C until used.

### Purification of recombinant NADP-ME variants

His-tagged proteins were purified using gravity-flow immobilized metal ion affinity chromatography on nickel-nitrilotriacetic acid (Ni-NTA) agarose. Frozen cells were thawed on ice and resuspended in 20 mM Tris-HCl (pH 8.0) containing 500 mM NaCl, 5 mM imidazole, 2 mM phenylmethanesulfonyl fluoride (PMSF), and a spatula-tip amount of lysozyme. The resuspended cells were sonicated for 8 min (30-s intervals with 30-s pauses) and then centrifuged at 15,000 × g for 15 min at 4 °C. The supernatant was loaded onto a pre-equilibrated Ni-NTA agarose column (Qiagen, Hilden, Germany) with a buffer containing 20 mM Tris-HCl (pH 8.0), 500 mM NaCl, and 5 mM imidazole. The column was subsequently washed with this buffer containing increasing concentrations of imidazole (5, 40, and 80 mM). Proteins were eluted with 2 ml of elution buffer containing 20 mM Tris-HCl (pH 8.0), 500 mM NaCl, and 500 mM imidazole. Finally, the eluted protein was concentrated and rebuffered with 20 mM Tris-HCl (pH 8.0) and 5mM MgCl_2_ at 4 °C using Amicon Ultra 0.5 ml Centrifugal Filters (50 kDa molecular weight cutoff) according to the manufacturer’s instructions (Merck, Darmstadt, Germany).

### Removal of His-tag

For His-tag removal, concentrated proteins were stored in the buffer recommended by the protease manufacturer. His-tags were cleaved using two different proteases, depending on the expression construct. For the pET16b vector, Factor Xa protease (New England Biolabs, Ipswich, USA) was used, while for the pET28b vector, Thrombin (Merck KGaA, Darmstadt, Germany) was employed. The recognition sites for both proteases are encoded in the respective expression vectors and are positioned between the His-tag and the NADP-ME sequence. The amount of protease used was in accordance with the manufacturer’s instructions, and cleavage was carried out at 4 °C for 16 hours. After cleavage, the His-tag and protease were removed by filtering the protein samples through Amicon Ultra 0.5 ml Centrifugal Filters (50 kDa molecular weight cutoff). The purified proteins were then rebuffered and stored in buffer containing 20 mM Tris-HCl (pH 8.0) and 5 mM MgCl_2_ at 4 °C, or in the same buffer with 20% glycerol at - 80 °C for long-term storage.

### Analytical Ultracentrifugation

Sedimentation velocity experiments were carried out using a Proteome Lab XL-A analytical ultracentrifuge (Beckman-Coulter, Brea, CA, USA). All recombinant versions of C4-NADP-ME and nonC4-NADP-ME were assayed in 20 mM Tris-HCl buffer (pH 7.0 or 8.0) with addition of 5 mM MgCl_2_. Final protein concentration was adjusted to 1.1 mg mL^−1^. Samples (16 µM) and corresponding buffer solutions were loaded into aluminium double sector centrepieces separately and built up in a Beckman An-60 Ti rotor. Experiments were performed at 20 °C and a rotor speed of 35,000 rpm. Protein samples were monitored by UV absorbance at λ = 280 nm in a continuous mode with a radial resolution of 0.003 cm. In time intervals of about 2 min scans of the radial concentration profile were collected until the protein was fully sedimented. Data were analyzed using the *c*(*s*) model in the software package SEDFIT (Schuck and Rossmanith, 2000). For data analysis, a resolution of 0.1 S with a confidence level (F-ratio) of 0.95 was chosen for the appropriate *s*-value range within 0 to 20.0 S. Density and viscosity of the solvent had been calculated with the software Sednterp (Laue *et al*., 1992) from tabulated values; ρ = 0.99823 g cm^−3^ and η = 0.01009 g cm^−1^s^−1^. The calculated partial specific volume of the individual proteins were approximately 0.738 cm³ g^−1^. Sedimentation coefficients are reported as *s*_20, w_ values, i.e. normalized to 20 °C and water as a solvent. Graphic output was generated by Gussi (Version 1.4.2) (Brautigam, 2015).

### Protein crystallization and X-ray data collection

For crystallization studies of C4G200R its His-tag was removed. The protein was crystallized using an Art Robbins Gryphon robot (Sunnyvale, CA, USA) on Jena Bioscience MRC 2-well sitting drop crystallization plates (Jena, Germany). Drops consisted of 0.2 µl of protein solution at 10 mg/ml in weak buffer (20 mM Tris-HCl pH 8.0, 5 mM MgCl_2_, 40 mM pyruvate, and 2 mM NADP) and 0.2 µl of commercial screen solution. Plates were incubated at 4 °C and were manually imaged on a Formulatrix Rock Imager 1000 apparatus (Bedford, MA, USA). C4G200R crystals appeared with a solution consisting of 2 M (MH_4_)SO_4_, 0.1 M Na acetate, pH 4.6. Samples were directly harvested from the robotic plates, cryo-protected in their respective mother liquors added with 35% (*w*/*v*) glycerol and flash cooled in liquid nitrogen for X-ray analysis.

### Data processing, structure determination and refinement

Native X-ray diffraction data were collected at the P13 beamline, operated by EMBL Hamburg at the PETRA III storage ring (DESY, Hamburg, Germany) (Cianci *et al*., 2017) using a Dectris EIGER X 16M detector (Baden, Switzerland). Indexing, integration and reduction was performed with XDS (https://xds.mr.mpg.de/) and Aimless-CCP4 (Agirre *et al*., 2023), leaving 5% of the reflections apart for cross-validation. The G200R structure (2.70 Å resolution) was solved by molecular replacement as implemented in CCP4 Cloud with MrBUMP (Keegan and Winn, 2008), using the coordinates of the wild-type enzyme (PDB code 5OU5) as starting model, followed by automatic model building with ModelCraft (Bond and Cowtan, 2022). Subsequent steps of manual model building and restrained refinement were carried out with Coot (Emsley *et al*., 2010) and Refmac5 (Murshudov *et al*., 2011), respectively. In the last cycles, a total of 6 pyruvate molecules and 29 water molecules were added to the model. The structure was validated with MolProbity (Chen *et al*., 2010) and with the validation module of Coot. Detailed information on data collection, refinement, validation and Protein Data Bank (PDB) deposition are shown in Suppl. Table 7.

### Cryo-EM sample preparation, data collection and processing

For cryo-EM, purified C4G200R and nonC4-NADP-ME without His-tag were prepared in 20 mM Tris-HCl, 5 mM MgCl_2_, 40 mM pyruvate, and 2 mM NADP (pH 8.0), and, purified C4G200R was additionally prepared in 10 mM Tris-HCl, 2.5 mM MgCl_2_, 20 mM pyruvate, 1 mM NADP, and 50 mM Na acetate (pH 4.8). To prepare the grids for cryo-EM, 4 μl of each sample solution (0.2-0.5 mg/ml) was applied to glow-discharged UltrAuFoil R1.2/1.3 grids (Quantifoil Micro Tools). The grids were blotted for 4.5 s and plunged frozen in liquid ethane after 0.5 s drain time in 100% relative humidity at 13 °C (Vitrobot Mark IV, Thermo Fisher Scientific). Automated datasets of the clipped grids were acquired using the EPU software with a Krios G4 Cryo-TEM (Thermo Fischer Scientific) at 300 kV, equipped with an E-CFEG, a Selectris X Energy Filter with a slit width of 10eV, and Falcon 4i Direct Electron Detector.

Each dataset (Suppl. Table 11) was pre-processed using CryoSPARC Live. The gain-corrected, motion-corrected, dose-weighted, and CTF-estimated micrographs were exported to CryoSPARC (v4.6.0) for further processing. Particles were initially blob-picked and subjected to several rounds of 2D classification. Classes containing contaminants were removed. The remaining particles were used to create templates to enhance the accuracy and efficiency of particle picking. The newly template-picked particles were subjected to new rounds of 2D classification in order to determine the oligomeric state of the samples. Suppl. Fig. 3 displays the processing workflow and all classes of the final round of 2D classification for each dataset, whereas Fig. 6C shows selected representative 2D class averages showing high-resolution features.

### Plant growth conditions and protein extract preparation

Maize seeds were soaked in water overnight, then wrapped in Whatman paper and secured inside the neck of a glass flask filled with water. The plants were grown at room temperature, either in the dark or under natural light conditions. After seven days, leaves and roots were harvested, rapidly frozen in liquid nitrogen, and ground into a fine powder. Protein extracts were then prepared following the protocol described by Hüdig *et al*. (2022).

### Protein quantification and gel electrophoresis

Protein concentration was determined using a modified amido black 10B precipitation method (Schaffner and Weissmann, 1973), with bovine serum albumin as the standard for the calibration curve, as described previously (Hüdig *et al*., 2022).

For native polyacrylamide gel electrophoresis (PAGE), proteins were separated on a non-denaturing 7% (*w/v*) polyacrylamide gel at 100 V and 4 °C. Protein bands were visualized either by Coomassie Brilliant Blue staining or through an in-gel NADP-ME activity assay.

### In-gel NADP-ME activity assay and immunological detection

NADP-ME activity was detected after native PAGE using an in-gel activity assay. The gels were first incubated in 50 mM Tris-HCl (pH 8.0) for 15 minutes at room temperature, followed by incubation in the NADP-ME activity assay solution, which contained 50 mM Tris-HCl (pH 8.0), 10 mM L-malate, 10 mM MgCl_2_, 0.5 mM NADP, 0.05% (*w/v*) nitro blue tetrazolium, and 150 µM phenazine methosulfate. After a brief wash with distilled deionized water (ddH_2_O), the gels were imaged.

For immunoblotting assays, the separated proteins were transferred to a PVDF membrane (Roti-PVDF, Carl Roth, Karlsruhe, Germany). After blocking with 5% milk powder in TBS, the membranes were incubated for at least 1 h with a 1:7,500 dilution of Anti-His-Tag antibody coupled to horseradish peroxidase (Anti-His-HRP, Miltenyi Biotec, Bergisch Gladbach, Germany). After washing several times with TBST, the membranes were incubated with a 1:2,500 dilution of Goat Anti-Rabbit IgG Antibody HRP-conjugate (Merck, Darmstadt, Germany). Proteins were visualized by chemiluminescence with Pierce ECL Western Blotting Substrate (Thermo Fisher Scientific, Waltham, USA) on a LAS-4000 Mini Luminescent Image Analyzer (GE Healthcare Life Sciences).

### Mass spectrometry

Protein extracts were separated on a 7% (*w/v*) native polyacrylamide gel electrophoresis (PAGE), and NADP-ME activity was visualized using an in-gel activity assay. The assay solution contained 50 mM Tris-HCl (pH 8.0), 100 mM L-malate, 10 mM MgCl_2_, 5 mM NADP, 0.05% (*w/v*) nitro blue tetrazolium, and 150 µM phenazine methosulfate. Gel slices exhibiting NADP-ME activity were excised and incubated for 20 min in a solution of 30% ethanol, 10% acetic acid, and 60% ddH_2_O. The gel slices were then prepared for mass spectrometric analysis, following the protocol described in Hüdig *et al*. (2022). Briefly, proteins were reduced with dithiothreitol (DTT), alkylated with iodoacetamide, and digested with trypsin. Tryptic peptides were extracted from the gel and purified using solid-phase extraction on an HLB µ-elution plate (Waters). Peptides were then separated using an Ultimate3000 rapid separation liquid chromatography system (Thermo Fisher Scientific, Waltham, USA) equipped with a C18 column, as previously described (Prescher *et al*., 2021). The peptides were injected into an online-coupled Fusion Lumos mass spectrometer (Thermo Fisher Scientific, Waltham, USA) with a FAIMS device via a nano electrospray source. The mass spectrometer was operated in data-dependent positive mode, with the FAIMS compensation voltage set to −50 V. Survey spectra were recorded in the Orbitrap analyzer (resolution 120,000, automatic gain control target 400,000, maximum ion time 60 ms, scan range 375-1500 m/z, profile mode). Precursor ions with 2-7 charges were isolated via the built-in quadrupole (isolation window 1.6 m/z), fragmented by higher-energy collisional dissociation (normalized collision energy: 35), and fragment spectra were recorded in the linear ion trap (scan rate: rapid, automatic gain control target 10,000, maximum ion time 150 ms, scan range: auto, centroid mode). The cycle time was set to 1 s, with dynamic exclusion of 1 min.

Mass spectrometry raw files were processed using Proteome Discoverer 2.4.1.15 for protein identification, based on the 63,255 protein sequences of the *Zea mays* reference proteome (UP000007305, downloaded from UniProt on October 19, 2023), as well as potential contaminants from MaxQuant (version 1.6.17.0, Max Planck Institute for Biochemistry, Planegg, Germany). The SequestHT search engine was used for the database search, with mass tolerances set to 10 ppm for precursor ions and 0.6 Da for fragment ions. Cysteine carbamidomethylation was set as a fixed modification, and methionine oxidation, N-terminal acetylation, and methionine loss were considered as variable modifications. Proteins and peptides were accepted at a 1% false discovery rate (FDR), validated based on peptide spectrum matches, and only proteins identified with at least two unique peptides were reported. Protein grouping was disabled, and all individual protein matches were reported.

To identify cytosolic NADP-ME, protein sequences from previously identified ones (Alvarez *et al*., 2013) were compared to the reference proteome, and the highest matching proteins were used for identification.

### Kinetic parameters and statistical analyses

NADP-ME activity, specifically the oxidative decarboxylation of L-malate, was determined spectrophotometrically by monitoring the formation of NADPH at 340 nm and 25 °C using a Tecan Spark plate reader (Tecan Group, Männedorf, Switzerland). The Michaelis constant (*K*_m_) for malate was determined at pH 8.0 by varying malate concentrations around the expected Km value, while keeping other components at saturating concentrations. The standard reaction mixture contained 50 mM Tris-HCl (pH 7.0 or 8.0), 10 mM MgCl_2_, 0.5 mM NADP, 200 ng of protein, and varying concentrations of L-malate ranging from 0.01 mM to 40 mM. For each assay, 192 µl of the reaction mixture was prepared in a Greiner 96-well flat transparent plate (Thermo Fisher Scientific, Waltham, USA). After recording the baseline measurement, 8 µl of increasing concentrations of L-malate were added to each well to achieve a final assay volume of 200 µl. NADPH formation was monitored at 340 nm and 25 °C for 25 min. Kinetic measurements were performed the day after protein purification, with proteins stored overnight at 4 °C. Activity measurements were repeated at least three times with independently expressed and purified protein samples, with each measurement consisting of triplicate assays for each concentration. Analysis of malate inhibition was performed at pH 7.0, with the remaining conditions staying the same. Kinetic parameters were calculated as described by Alvarez *et al*. (2019), using data from three independently purified protein batches and triplicate measurements for each. The extinction coefficient (ε) of 6.22 mM^−1^ cm^−1^ at 340 nm for NADPH was used in the calculations. Data were fitted to non-linear regression in GraphPad Prism version 8.3.0 software (GraphPad Software, Boston, Massachusetts, USA) using free concentrations of all substrates.

Statistical analysis of kinetic measurements was performed with GraphPad Prism 8.3.0 software using a two-tailed t-test with Welch’s correction (α < 0.05), comparing nonC4- and C4-NADP-ME variants to the corresponding wild-type proteins. P-values are shown in Suppl. Table 2.

### Phylogenetic Analysis

Multiple sequence alignment was performed as described in Tronconi *et al*. (2020). Maximum likelihood analysis of the complete dataset was carried out using MEGA X (v.10.0.5). The fit of each model to the data was evaluated using the Bayesian Information Criterion, with the model yielding the lowest BIC score being selected as the most appropriate for describing the substitution pattern. The initial ML tree was generated automatically using the NJ and BIONJ algorithms, and branch lengths were optimized to maximize the likelihood of the dataset under the chosen evolutionary model. Heuristic searches were then conducted starting from this initial tree, employing the nearest neighbor method, where alternative trees differ in one branching pattern. The reliability of internal branches was assessed through 2,000 bootstrap resamplings. Bootstrap values of 50-69% were considered weakly supported, 70-84% as moderately supported, and 85-100% as strongly supported (Hillis and Bull, 1993). The tree was saved in Newick format (.nwk), containing all relevant clade support values and branch length information, and visualized using FigTree v1.4.4 software.

### Analysis of interactions and residues involved in oligomerization

Analyses of hydrogen bonds (Suppl. Table 4) and salt bridges (Suppl. Table 5) between monomers of a dimer and identification of residues with stabilizing or destabilizing roles in oligomer formation (Suppl. Table 6) were performed using the PISA program (CCP4; (Krissinel and Henrick, 2007). Critical residues involved in stabilizing the oligomeric structure were identified through HOT Spot analysis using the KFC2 program (Zhu and Mitchell, 2011) (Suppl. Table 3). Hydrogen bond analysis and quality figures were generated with the licensed PyMOL Molecular Graphics System, Version 2.4.0 Schrödinger, LLC.

### Data availability

The data supporting the findings of this manuscript are available from the corresponding author upon reasonable request. The C4G200R crystallographic structure was deposited in the Protein Data Bank (PDB) under the accession code PDB ID 9E6M.

## Acknowledgements

This work was supported by the Deutsche Forschungsgemeinschaft (DFG) grant MA2379/20-1 (project number 441941117) awarded to VGM and the National Agency for Promotion of Science and Technology (ANPCyT) grant PICT-2019-00079 to CEA. We extend our heartfelt gratitude to Nils Schneberger and Matthias Geyer from the University Clinics Bonn, University of Bonn, Germany, for their assistance with the crystallographic facility, and to Julie Mitchell from the University of Wisconsin-Madison, USA, for her valuable support with Hot Spots analysis.

The authors declare that they have no conflicts of interest.

