## Supplementary material for "From dimer to tetramer: the evolutionary trajectory of C4 photosynthetic-NADP-ME oligomeric state in Poaceae"

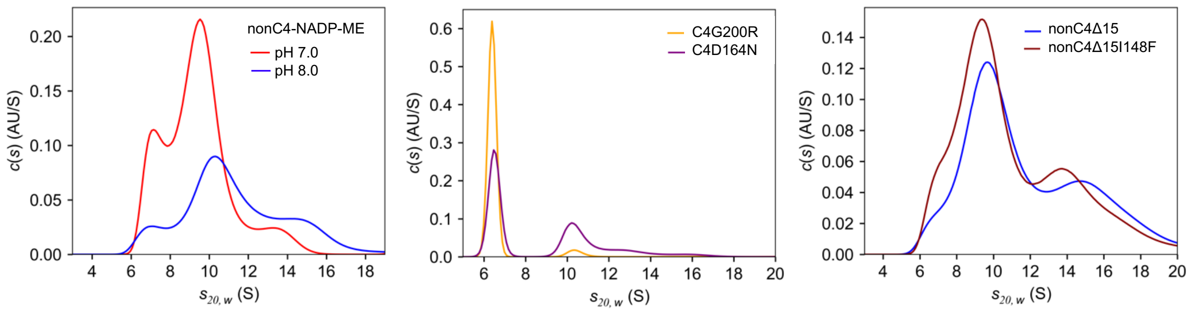


**Supplementary Figure 1. Continuous sedimentation coefficient distribution of C4- and non-C4 NADP-ME variants.** Unless otherwise specified, measurements were conducted at pH 8.0. Data were analyzed using the c(s) model in the software package SEDFIT.

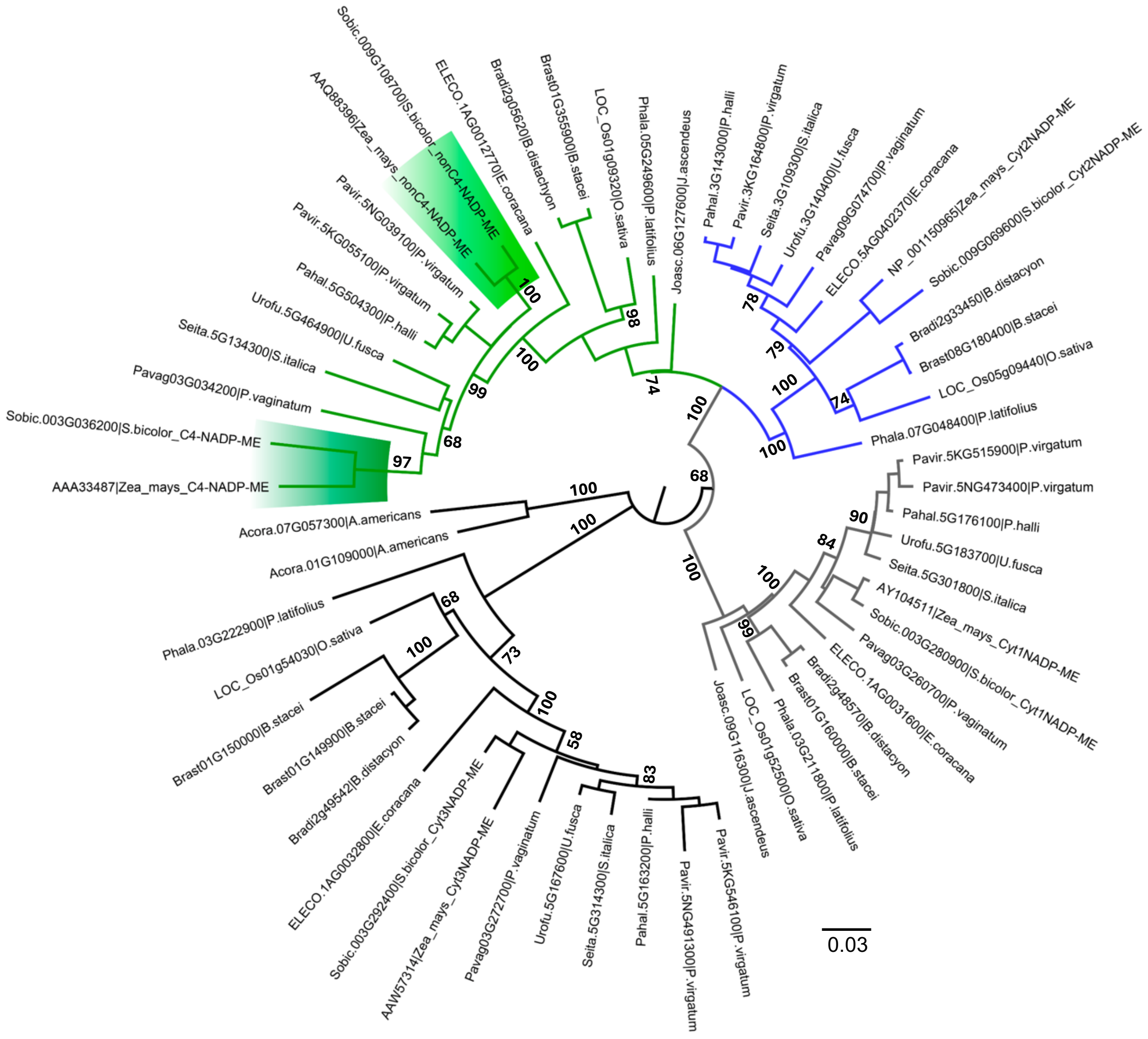

**Supplementary Figure 2. Phylogenetic tree of NADP-ME proteins in monocot.** The evolutionary relationships were inferred using the Maximum Likelihood (ML) method, based on a Multiple Sequence Alignment of 58 protein sequences spanning 554 amino acid positions from the genomes of 18 monocot species. *Acorus americanus*, an early-branching monocot, was used as the outgroup. The tree is drawn to scale, with branch lengths representing the number of substitutions per site. Evolutionary distances were calculated using the JTT matrix-based model, with a discrete Gamma distribution (parameter=0.52) to account for rate variation among sites. Bootstrap support values (from 2,000 replicates) are shown as MLB values next to branches, with values greater than 50% indicated. Cytosolic lineages I, II, and III are represented in black, gray, and blue, respectively. The plastidic lineage is shown in green. Maize and sorghum C4-NADP-ME are highlighted in dark green, while non-C4-NADP-ME are indicated in light green.

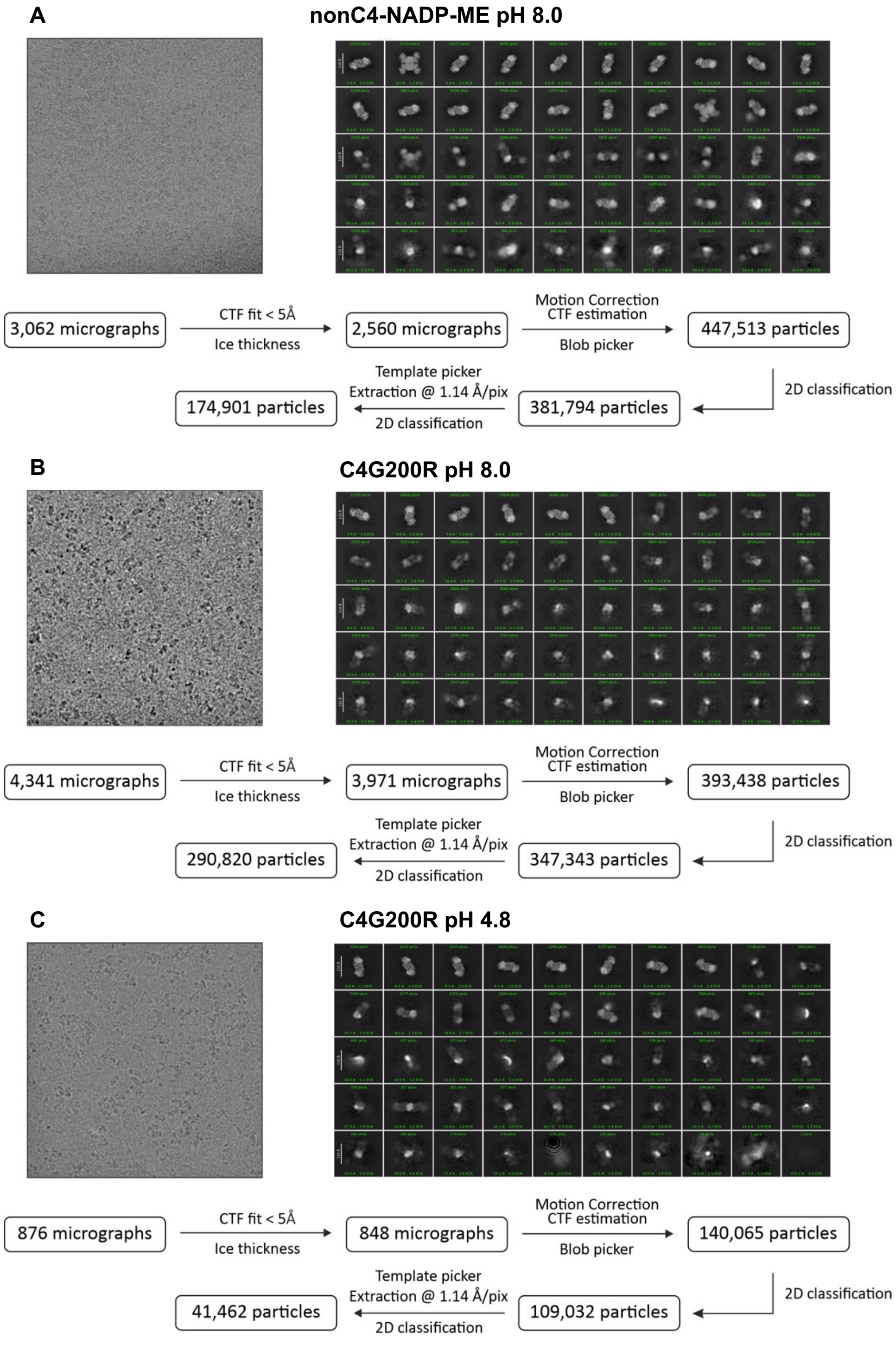


**Supplementary Figure 3. Cryo-EM analysis workflow of nonC4-NADP-ME and C4G200R.** A representative micrograph, 2D class averages and flowchart for cryo-EM data processing are shown for nonC4-NADP-ME, pH 8.0 (upper panel), C4-NADP-ME-G200R, pH 8.0 (middle panel) and C4G200R, pH 4.8 (lower panel).

```
AtNAD-ME1           ------------------------------------------------------------------------ATGGGAATAGCCAATAAGCTCCGGCTA
AtNAD-ME2           ------------------------------------------------------------------------------------------------ATG
GgNAD-ME1           -----------------------------------------------------------------------------ATGGCGTCTGTCGGAGATAAGC
GgNAD-ME3           ---------------------------------------------------------------------------------------------------
ZmC4NADP-ME         ---------ATGCTGTCCACGCGCACCGCCGCCGTCGCCGCCTCCGCGTCCCCCGCCTCCCCGTGGAAGCTGGGAGGACGGAGCGAGGGCGGCGCCAGC
SbC4NADP-ME         ---------ATGATCTCCGCTCGCGCCGCCACCGTCGCCGCCGCGTCCCCCGCCTCCCCGTGGAAGCGGGGAGGAGGACGGAGCGAGGGTGCCGGCAGC

ZmnnonC4-NADP-ME    ------------------------ATGATCTCCGCGCGCACCGCCGCAGCTTCCCCCGCTTCCCTGTGGAAGCAGGGAGGACGAAGCGAGGGCGGCGGC
SbnonC4-NADP-ME     ------ATGATCTCCGCGCGCACCGCCGCGTCCCCGGCCTCCCCCGCCTCCGCGTGGAAGCGGGGAGGAGGACGAAGCGAGGTCGGCGGCGGCAGCAGC


                                                                                         a       b   c  d   e
AtNAD-ME1           AGTTCATCATCTCTCAGCCGAATCCTCCACCGGAGAATACTTTACTCATCCGCTGTCAGATCTTTCTTCACCACATCGGAAGGTCACCGTCCCACCATC
AtNAD-ME2           ATGTGGAAGAACATTGCTGGGTTGTCGAAGGCAGCGGCAGCGGCAAGAACACACGGATCTCGGCGGTGCTTTTCCACAGCGATTCCTGGTCCTTCCATC
GgNAD-ME1           TCCGGATTCTCGAGGGCTATGACTATGATTCTTAGACGTAGGATTTTGGGTGAAACCCCCAGACGTTTCACTACAACGGAGGGCCACCGTCCCACCATT
GgNAD-ME3           ---ATGATTTGGAGGACTGTTGCCAGATTTGCGACGTCGGCGGCGAGAGCCGGCGGATCTCATCGATGTTTTTCGACGGCGATTCCAGGACCTTCCATT
ZmC4-NADP-ME        TGCGACGGATGCCGGACCTACAGGAACACCCTGCGGAGGAGGGCCGCGCCCGCCAAGGTGCGCGCCTTGCCGCCCAGGCGGGTGGACGCGGTCGCGATG
SbC4-NADP-ME        TGCGACGGATGCAGCACCTACAGGAAGACCGTGCGGAGGAGGGCCGCCGCCGCCAAGGTGCGCGCCTTGCCGCCGAAGCGGGTGGAGGCGGTCGCGATA

ZmnnonC4-NADP-ME    AGCTGCGACGGATGTCGGACTTACAGGAACACTTTGCGGAGGGCCGCAGCAGCGAAGGTGCGTGCCTTGCCGTCGAGGGCGGTGGAGGCGGTCAAAATG
SbnonC4-NADP-ME     TGCGACGGATGCCGGACCTACAGGAACACCTTGCGGAGGAGCGCCGCGGCGGCGAAGGTGCGCGCCGTGCCGCCGAGGCGGGTGGAGGCGGTCGCGATG


                    f     g                      BOX 1               h  i   j   k   l   m  n           o  p q
AtNAD-ME1           GTTCATAAACAAGGTCTCGATATCCTCCATGATCCTTGGTTCAACAAGGGGACTGCGTTTACGATGACGGAGAGAAATCGTTTAGATCTTAGAGGTCTT
AtNAD-ME2           GTCCACAAGCGTGGTGCTGATATTCTTCACGATCCATGGTTCAATAAGGACACAGGTTTTCCTTTGACTGAGAGAGATAGATTAGGGATACGAGGCTTG
GgNAD-ME1           GTCCACAAGCGAAGCCTCAACATCCTTCACGATCCTTGGTTCAACAAGGGAACTGCCTTCTCAATGACTGAGAGAGACCGTCTTGATCTCAGGGGTCTT
GgNAD-ME3           GTCCATAAGCGTGGCGCTGATATTCTTCATGATCCCTGGTTCAATAAGGATACAGGTTTTCCTTTGACTGAGAGAGACAGATTAGGGCTCCGAGGCCTT
ZmC4NADP-ME         GGCTCCGACGCGGAGACGGAGACGGAGAAGGAGCAGGAGGAGGCGGCGGCCGCCAGCGAGGAGCTGCCCGTCATGCCCTGGGCCACCTCCGTCGCAAGC
SbC4NADP-ME         GGCTCCAGCGCGGAGACGGAGACGGAGGAGGTGGTGGTGGAGGTGGCGGCGGCCAACGAGGAGCTGCCCGTCATGCCCTGGGCCACCTCTGTCGCAAGC
                                                                                    a       b   c  d   e    f   g
ZmnnonC4-NADP-ME    GGCTCCGCCGCGGAGATGGAGCAGGAGGAGGAGGAGGCGGAGGTGGCCGCGGCCGCGGCCGGCGTCGTCGTCGAGGACCACTACGGCGAGGACAGCGCC
SbnonC4-NADP-ME     GGCTCCGCCGCGGAGACGGAGAAGGAGAAGGAGGAGCAGCAGGAGGTGGTGGCCGGGGCCGGCGGCCTCGTCGAGGACCACTACGACGAGGACAGTGCC
Cyt2                                                                ATGGCGGGCGGCGGCGTTGAGGACGTGTACGGCGAGGACCGCGCT


                      BOX 2      r  s
AtNAD-ME1           CTTCCTCCTAATGTTATGGACTCTGAGCAACAGATTTTTCGTTTTATGACTGACCTGAAGAGATTAGAGGAGCAAGCTAGAGATGGACCTTCTGATCCT
AtNAD-ME2           CTTCCTCCTCGTGTTATGACCTGTGTACAGCAATGTGATCGTTTCATTGAGTCGTTTCGTTCGCTAGAGAACAACACAAAAGGAGAGCCTGAAAACGTT
GgNAD-ME1           CTTCCTCCGAACGTCATGGCTCCTGAGCAGCAAATCGAACGCTTCATGGCGGACTTTAAGAGGCTTCAGAAGCAAGCAAGGGATGGACCGCTTGATCCT
GgNAD-ME3           CTTCCTCCTCGTGTAATCTCATTCGAACAGCAATATACCCGTTTCATGGAATCTTTTCGATCACTAGAGAGAAATACTAAGGGACAGCCAGAAGATGCT
ZmC4NADP-ME         GGTTACACCCTTTTGAGGGATCCACACCACAACAAGGGTCTTGCTTTCACAGAGGAGGAAAGGGATGGACACTACTTGCGTGGACTGCTTCCTCCGGCA
SbC4NADP-ME         GGTTACACCCTTTTGAGGGATCCACGTCACAACAAGGGTCTTGCTTTCACGGAGGAAGGAGAGGGATGCACACTACTTGCGTGGACTGCTTCCTCCGGCA
                                          BOX 1              h   i    j   k    l    m  n            o   p q     BOX 2
ZmnnonC4-NADP-ME    ACCGAGGAGCTGCCCATTTTGCCCTGGGCCTTCTCCGTCGCAAGCGGTTACACCCTCTTGAGGGATCCACATCACAACAAGGGTCTTGCTTTCATGGAG
SbnonC4-NADP-ME     GCCGAGGAGGTGCCAATCATGCCCTGGGCCTTCTCCGTCGCGAGCGGTTACACTCTCTTGAGGGATCCCCATCACAACAAGGGTCTTGCTTTCACGGAG
Cyt2                ACCGAGGAGCAGCTGATCACGCCGTGGGCCTTCTCCGTCGCAAGCGGCTACACCCTGCTGCGCGACCCGCGGCACAACAAGGGCCTGGCCTTCTCGGAG
                                  a       b   c  d    e    f     g                            BOX 1              h   i   j  k


AtNAD-ME1           AATGCTTTGGCTAAATGGCGGATTCTTAATCGGTTACATGACCGAAATGAGACTATGTACTATAAGGTTTTGATTAACAATATTGAGGAGTATGCGCCG
AtNAD-ME2           GTTGCATTAGCTAAATGGAGGATGTTGAATAGATTGCATGACCGTAATGAGACTTTGTACTACCGAGTTCTTATTGATAACATCAAAGATTTTGCCCCC
GgNAD-ME1           AATGCTCTGGCTAAGTGGCGTATCCTTAATCGTCTGCATGACCGAAACGAGACCATGTACTTTAAGGTTTTGATTGCCAATATCGAGGAATATGCGCCT
GgNAD-ME3           GTATCATTAGCAAAATGGAGGATATTGAATCGATTACATGATCGTAACGAGACTTTGTATTACCGAGTCCTTATTGATAACATCAAAGACTTTGCTCCA
ZmC4NADP-ME         GTACTATCTCAGGAACTCCAAATTAAGAAGTTCATGAACACCCTGCGGCAGTACCAGACCCCTTTGCAGCGGTATATAGCTATGATGAACCTTCAGGAA
SbC4NADP-ME         GTTGTCTCTCAGGAACTCCAAATTAAGAAGTTCATGAACAACCTGCGGCAGTACCAGCTCCCTATTCAGTGCTATATGGCCATGATGAACCTTCAGGAG
                     r  s
ZmnnonC4-NADP-ME    AAGGAGAGGGATGCACACCACTTGCGGGGCTGCTTCCTCCTGCAGTTGTGCCTCAAGAACTCCAAATTAAGAAGATCATGCACAACCTGCGGCAGTAC
SbnonC4-NADP-ME     AAGGAGAGGGATGCACACCACTTGCGTGGCTGCTTCCTCCTGCAGTTGTGCCTCAGCAACTCCAAGTTAAGAAGATCATGCACAACCTGCGGCAGTAC
Cyt2                GCGGAGCGCGACGCGCACTACCTCCGGGGCCTGCTGCCGCCGGCGCTGGCGTCGCAGGAGCTGCAGGAGAAGAAGCTCATGCAGAACCTGCGCCGGTAC
                       l   m  n          o  p q     BOX 2      r  s
```

**Supplementary Figure 4. Alignment of 5´coding sequences showing regions of identical nucleotide sequences shared between monocot and dicot NAD(P)-ME enzymes.** Predicted transit peptide cleavage sites are shown in bold. The sequence encoding the additional residues found in the N-terminal region of maize and sorghum nonC4-NADP-ME are shown in orange. Elements (a to s) and Box 1 and Box 2 are annotated according to Brown *et al.* (2011).

**Supplementary Table 1. Residues strictly differentially substituted between C4- and nonC4-NADP-ME in maize and sorghum as described in Alvarez *et al.* (2019).** Amino acid numbering corresponds to positions in the full-length sequence of maize C4-NADP-ME, and the positional homologous residues of the full-length nonC4-NADP-ME.

| C4-NADP-ME | nonC4-NADP-ME |
|---|---|
| T92 | F100 |
| F140 | I148 |
| N142 | H150 |
| N159 | D167 |
| T163 | R171 |
| D164 | N172 |
| V177 | E185 |
| G200 | R208 |
| R201 | Q209 |
| D266 | R274 |
| D304 | H312 |
| F317 | I325 |
| E339 | A347 |
| M369 | V377 |
| Q377 | H385 |
| I474 | V482 |
| Q503 | E511 |
| T506 | N514 |
| A521 | D529 |
| L544 | F552 |

**Supplementary Table 2. P-values from the kinetic parameters shown in Table 1.** P-values from two-tailed t-tests with Welch's correction, comparing the kinetic parameters of nonC4-NADP-ME variants to the nonC4 wild-type enzyme and C4-NADP-ME variants to the C4 wild-type isoform. (*) P < 0.05; (**) P < 0.01; (***) P < 0.001. (ns) not statistically significant.

| NADP-ME | | P-value $K_m$ | P-value $k_{cat}$ |
|---|---|---|---|
| nonC4 | DelN | 0.2927 (ns) | 0.2180 (ns) |
| | Δ15 | 0.0933 (ns) | 0.0549 (ns) |
| | Δ15I148F | 0.9056 (ns) | 0.0326 (*) |
| | Δ15IF_AE | 0.0152 (*) | 0.0005 (***) |
| | Δ15IF_AE_EQ | 0.0271 (*) | 0.0062 (**) |
| | Δ15IF_AE_FL | 0.0131 (*) | 0.1591 (ns) |
| | Δ15IF_AE_EQ_FL | 0.0830 (ns) | 0.0926 (ns) |
| | Δ15_20aa | 0.0101 (*) | 0.7492 (ns) |
| | R171T | 0.0040 (**) | 0.0029 (**) |
| | N172D | 0.7575 (ns) | <0.0001 (***) |
| | R208G | 0.4025 (ns) | <0.0001 (***) |
| C4 | G200R | 0.0126 (*) | <0.0001 (**) |
| | +15 | 0.6552 (ns) | <0.0001 (***) |
| | NnC4 | 0.7574 (ns) | <0.0001 (***) |

**Supplementary Table 3. Residues identified as hot spots (HS) using the KFC2 machine learning program.** HS exclusive for each protein are highlighted in bold. An asterisk, indicates the HS introduced by the G200R mutation in C4-NADP-ME (C4G200R).

| | **nonC4-NADP-ME** | **C4-NADP-ME** | **C4G200R** |
|---|---|---|---|
| Leu | 109 | 101 | 101 |
| Arg | 110 | 102 | 102 |
| Leu | 117 | 110 | 110 |
| Arg | 131 | 123 | 123 |
| Gly | 132 | 124 | 124 |
| Leu | 133 | 125 | 125 |
| Leu | 134 | 126 | 126 |
| Pro | 135 | 127 | 127 |
| Pro | 136 | 128 | 128 |
| Lys | **147** | - | **139*** |
| Tyr | **155** | - | **147*** |
| Asn | **167** | - | **159*** |
| Arg | **171** | - | **163*** |
| Phe | 207 | 199 | 199 |
| Arg | **208** | - | **Arg 200*** |
| Gln | - | **Arg 201** | **Arg 201** |
| Pro | 210 | 202 | 202 |
| Leu | 213 | 205 | 205 |
| Tyr | 214 | 206 | 206 |
| Asn | 229 | 221 | 221 |
| Ser | - | **Cys 246** | **246** |
| Phe | 297 | 289 | 289 |
| Tyr | 298 | 290 | 290 |
| Ile | 199 | 291 | 291 |

**Supplementary Table 4. Hydrogen bond interactions between chains A and B in C4- and nonC4-NADP-ME, identified by PISA analysis.** The table details the residues and atoms participating in the interactions, along with the corresponding distances (Å). Isoform-specific differences are color-coded: green for C4-NADP-ME and orange for nonC4-NADP-ME. Residues labeled with ʃ indicate amino acid differences between the isoforms, while residues marked with * belong to the loop containing G200 in C4-NADP-ME.

| Residue (chain A) | Atom | Distance (Å) | Residue (chain B) | Atom | Equivalent residue to: | |
|---|---|---|---|---|---|---|
| **C4-NADP-ME** | | | | | **nonC4-NADP-ME** | |
| P202(*) | $O^{\eta}$ | 3.6 | Y98 | $O^{\eta}$ | P210(*) | Y106 |
| **I291** | **O** | **3.1** | **A129** | **N** | **I299** | **A137** |
| G196 | O | 2.8 | **K139** | $N^{\zeta}$ | G204 | **K147** |
| **Q146** | **O** | **3.2** | **Q152** | $N^{\varepsilon 2}$ | Q154 | **Q160** |
| N159(ʃ) | $O^{\delta 1}$ | 3.5 | N159(ʃ) | $N^{\delta 1}$ | D167(ʃ) | D167(ʃ) |
| N159(ʃ) | $N^{\delta 2}$ | 3.4 | N159(ʃ) | $O^{\delta 1}$ | D167(ʃ) | D167(ʃ) |
| L101 | O | 3.4 | R201(*)(ʃ) | $N^{\eta 1}$ | L109 | Q209(*) (ʃ) |
| L101 | O | 3.7 | R201(*)(ʃ) | $N^{\eta 2}$ | L109 | Q209(*) (ʃ) |
| **G124** | **O** | **2.8** | **Y206** | **N** | G132 | Y214 |
| **R123** | **O** | **3.0** | **N221** | $N^{\delta 2}$ | R131 | N229 |
| **N221** | $N^{\delta 1}$ | **2.9** | **R123** | **O** | N229 | R131 |
| **Y206** | **N** | **2.8** | **G124** | **O** | Y214 | G132 |
| **Q152** | $N^{\varepsilon 2}$ | **3.2** | **Q146** | **O** | Q160 | **Q154** |
| **Q152** | $N^{\varepsilon 2}$ | **3.5** | Q148 | $O^{\varepsilon 1}$ | Q160 | Q156 |
| **K139** | $N^{\zeta}$ | **2.8** | G196(*) | O | K147 | G204 |
| Y98 | $O^{\eta}$ | 3.4 | P202(*) | O | Y106 | P210(*) |
| **A129** | **N** | **3.1** | **I291** | **O** | A137 | I299 |
| **nonC4-NADP-ME** | | | | | **C4-NADP-ME** | |
| R110 | $N^{\eta 1}$ | 3.1 | R110 | O | R102 | R102 |
| **N229** | $N^{\delta 2}$ | **2.8** | **R131** | **O** | N221 | R123 |
| **Y214** | **N** | **2.6** | **G132** | **O** | Y206 | G124 |
| R218(ʃ) | $N^{\eta 1}$ | 3.8 | P136 | O | K210(ʃ) | P128 |
| **Q160** | $N^{\varepsilon 2}$ | **3.7** | **Q154** | **O** | Q152 | Q146 |
| R208(*)(ʃ) | $N^{\eta 2}$ | 3.7 | Y155 | $O^{\eta}$ | G200(*)(ʃ) | Y147 |
| R110 | $N^{\eta 2}$ | 3.2 | E170 | O | R102 | E162 |
| N151(ʃ) | $N^{\delta 2}$ | 2.8 | S205 | O | T143(ʃ) | S197 |
| **K147** | $N^{\zeta}$ | **2.9** | F207(*) | O | K139 | F199 |
| **R131** | $N^{\eta 1}$ | **3.9** | R218(ʃ) | O | R123 | K210(ʃ) |
| **R131** | **O** | **2.9** | **N229** | $N^{\delta 2}$ | R123 | N221 |
| **A137** | **N** | **3.1** | **I299** | **O** | A129 | I291 |
| R110 | O | 3.2 | R110 | $N^{\eta 1}$ | R102 | R102 |
| **G132** | **O** | **2.6** | **Y214** | **N** | G124 | Y206 |
| P136 | O | 3.8 | R218(ʃ) | $N^{\eta 1}$ | P128 | K210(ʃ) |
| **Q154** | **O** | **3.7** | **Q160** | $N^{\varepsilon 2}$ | Q146 | Q152 |
| Y155 | $O^{\eta}$ | 3.7 | R208(*)(ʃ) | $N^{\eta 2}$ | Y147 | G200((*)(ʃ) |
| E170 | O | 3.3 | R110 | $N^{\eta 2}$ | E162 | R102 |
| S205 | O | 2.8 | N151(ʃ) | $N^{\delta 2}$ | S197 | T143(ʃ) |
| F207(*) | O | 2.9 | **K147** | $N^{\zeta}$ | F199 | K139 |
| R218(ʃ) | O | 3.8 | **R131** | $N^{\eta 1}$ | K210(ʃ) | R123 |
| **I299** | **O** | **3.1** | **A137** | **N** | I291 | A129 |

**Supplementary Table 5. Salt bridge interactions between monomers A and B for C4- and nonC4-NADP-ME, identified by PISA analysis.** The table lists the interacting atoms and the corresponding distances (Å) for each residue involved. Conserved interactions shared by both isoforms are highlighted in bold, while isoform-specific atoms participating in these interactions are marked in orange. Residues labeled with (¥) are located at the C-terminus of the αA1 helix, and those marked with (£) are situated within the αA3 helices in both isoforms.

| Residue (chain A) | Atom | Distance (Å) | Residue (chain B) | Atom | Equivalent residue to | |
|---|---|---|---|---|---|---|
| C4-NADP-ME | | | | | nonC4-NADP-ME | |
| **D211** | **$O^{\delta1}$** | **3.9** | **R123** | $N^{\varepsilon}$ | D219 | R131 |
| **E288** | **$O^{\varepsilon1}$** | **2.8** | **K138(£)** | **$N^{\zeta}$** | E296 | K146 |
| **E288** | **$O^{\varepsilon2}$** | **3.6** | **K138(£)** | **$N^{\zeta}$** | E296 | K146 |
| **R102(¥)** | **$N^{\eta2}$** | **3.7** | **E162** | **$O^{\varepsilon2}$** | R110 | E170 |
| **K138(£)** | **$N^{\zeta}$** | **2.8** | **E288** | **$O^{\varepsilon1}$** | K146 | E296 |
| **K138(£)** | **$N^{\zeta}$** | **3.5** | **E288** | **$O^{\varepsilon2}$** | K146 | E296 |
| nonC4-NADP-ME | | | | | C4-NADP-ME | |
| **R110(¥)** | **$N^{\eta2}$** | **2.7** | **E170** | **$O^{\varepsilon2}$** | R102 | E162 |
| **R131** | $N^{\eta1}$ | 3.5 | **D219** | $O^{\delta1}$ | E123 | D211 |
| **K146(£)** | **$N^{\zeta}$** | **3.8** | **E296** | **$O^{\varepsilon1}$** | K138 | E288 |
| **K146(£)** | **$N^{\zeta}$** | **3.7** | **E296** | **$O^{\varepsilon2}$** | K138 | E288 |
| **E170** | **$O^{\varepsilon2}$** | 2.7 | **R110(¥)** | $N^{\eta2}$ | E162 | R102 |
| **D219** | **$O^{\delta1}$** | **3.4** | **R131** | $N^{\eta1}$ | D211 | R123 |
| **E296** | **$O^{\varepsilon1}$** | **3.8** | **K146(£)** | **$N^{\zeta}$** | E288 | K138 |
| **E296** | **$O^{\varepsilon2}$** | **3.6** | **K146(£)** | **$N^{\zeta}$** | E288 | K138 |

**Supplementary Table 6. Stabilizing and destabilizing residues involved in the dimer interface of C4- and nonC4-NADP-ME, identified by PISA analysis.** The analysis includes type of residues involved. The residues T163, R201, and P202 are highlighted in bold (see the main text for details).

| Stabilizing residues between monomers contacts | | Equivalent residue in |
|---|---|---|
| | Residue | nonC4-NADP-ME |
| C4-NADP-ME | F289 | F297 |
| | **P202** | P210 |
| | P128 | P136 |
| | P127 | P135 |
| | A129 | A137 |
| | Y206 | Y214 |
| | | C4-NADP-ME |
| nonC4-NADP-ME | P136 | P128 |
| Destabilizing residues between monomers contacts | | |
| | | nonC4-NADP-ME |
| C4-NADP-ME | **R201** | **Q209** |
| | | C4-NADP-ME |
| nonC4-NADP-ME | R171 | **T163** |

**Supplementary Table 7. X-ray data collection and refinement statistics for the crystal structure of C4G200R.** [a] Statistics for the highest resolution shell are given in parentheses. [b] RMS: Root-mean square.

| | |
|---|---|
| **Data collection:** | |
| Crystal-detector distance (mm) | 407.3 |
| Rotation range/image (°) | 0.2 |
| No. of frames | 700 |
| Exposure time/image (s) | 0.021 |
| Wavelength (Å) | 0.9762 |
| Space group | $P2_12_12_1$ |
| Unit cell parameters | |
| *a*, *b*, *c* (Å) | 98.68, 124.16, 189.47 |
| *α*, *β*, *γ* (°) | 90, 90, 90 |
| Resolution range (Å) [a] | 50.00–2.70 (2.77–2.70) |
| Total reflections | 341,332 (24,753) |
| Unique reflections | 63,889 (4,473) |
| Redundancy | 5.3 (5.5) |
| Completeness (%) | 99.1 (99.8) |
| Mean I/σ(I) | 5.3 (1.1) |
| Overall Wilson *B*-factor (Å$^2$) | 50 |
| $R_{meas}$ | 0.213 (1.865) |
| $R_{pim}$ | 0.092 (0.780) |
| $CC_{(1/2)}$ | 0.993 (0.620) |
| Subunits/asymmetric unit | 4 |
| **Refinement:** | |
| Reflections used in refinement | 60,592 |
| $R_{free}$ test set (%) | 5.0 |
| $R_{work}$ | 0.239 |
| $R_{free}$ | 0.296 |
| No. of non-hydrogen atoms | |
| all atoms | 17,498 |
| macromolecules | 17,433 |
| ligands | 36 |
| solvent | 29 |
| RMS [b] deviations from ideal values | |
| bonds (Å) | 0.010 |
| angles (°) | 1.71 |
| Average *B*-factor (Å$^2$) | |
| all atoms | 66 |
| protein | 66 |
| ligand | 83 |
| solvent | 35 |
| Ramachandran plot | |
| favored regions (%) | 94.5 |
| allowed regions (%) | 5.1 |
| outliers (%) | 0.4 |
| **Deposition:** | |
| PDB code | 9E6M |

**Supplementary Table 8. Hydrogen bond interactions between chains A and B in C4- and C4G200R, identified by PISA analysis.** The table details the residues and atoms participating in the interactions, along with the corresponding distances (Å). Isoform-specific differences are color-coded: green for C4-NADP-ME and yellow for C4G200R. Residues labeled with * belong to the loop containing G200 in C4-NADP-ME.

| Residue (chain A) | Atom | Distance (Å) | Residue (chain B) | Atom | Residue (chain A) | Atom | Distance (Å) | Residue (chain B) | Atom |
|---|---|---|---|---|---|---|---|---|---|
| C4-NADP-ME | | | | | G200R | | | | |
| P202(*) | O | 3.6 | Y98 | $O^{\eta}$ | R123 | $N^{\eta1}$ | 3.2 | K210 | O |
| **I291** | **O** | 3.1 | **A129** | **N** | **A129** | **N** | 3.2 | **I291** | **O** |
| **G196** | **O** | 2.8 | **K139** | $N^{\zeta}$ | **Q148** | $N^{\varepsilon1}$ | 3.6 | **Q152** | $O^{\varepsilon2}$ |
| **Q146** | **O** | 3.2 | **Q152** | $N^{\varepsilon2}$ | **Q152** | $N^{\varepsilon2}$ | 3.3 | **Q146** | **O** |
| **N159** | $O^{\delta1}$ | 3.5 | N159 | $N^{\delta1}$ | **Y206** | **N** | 2.8 | **G124** | **O** |
| **N159** | $N^{\delta2}$ | 3.4 | **N159** | $O^{\delta2}$ | N221 | $N^{\delta2}$ | 2.7 | **R123** | **O** |
| **L101** | **O** | 3.4 | **R201(*)** | $N^{\eta1}$ | K210 | O | 3.3 | R123 | $N^{\eta1}$ |
| L101 | O | 3.7 | R201(*) | $N^{\eta2}$ | **I291** | **O** | 3.1 | **A129** | **N** |
| **G124** | **O** | 2.8 | **Y206** | **N** | **G196** | **O** | 2.8 | **K139** | $N^{\zeta}$ |
| **R123** | **O** | 3.0 | **N221** | $N^{\delta2}$ | **Q146** | **O** | 3.2 | **Q152** | $N^{\varepsilon2}$ |
| **N221** | $N^{\delta2}$ | 2.9 | **R123** | **O** | **N159** | $O^{\delta1}$ | 3.4 | **N159** | $N^{\delta2}$ |
| **Y206** | **N** | 2.8 | **G124** | **O** | **L101** | **O** | 3.8 | **R201(*)** | $N^{\eta1}$ |
| **Q152** | $N^{\varepsilon2}$ | 3.2 | **Q146** | **O** | **G124** | **O** | 2.7 | **Y206** | **N** |
| **Q152** | $N^{\varepsilon2}$ | 3.5 | Q148 | $O^{\varepsilon1}$ | **R123** | **O** | 2.6 | **N221** | $N^{\delta2}$ |
| **K139** | $N^{\zeta}$ | 2.8 | **G196** | **O** | | | | | |
| Y98 | $O^{\eta}$ | 3.4 | P202(*) | O | | | | | |
| **A129** | **N** | 3.1 | **I291** | **O** | | | | | |

**Supplementary Table 9. List of vectors used for recombinant protein expression.** The table includes the name of the encoded protein, the corresponding vector backbone, the method of vector construction (Gene synthesis by BioCat, SDM: site directed mutagenesis, or Gibson cloning), and the *E. coli* strain used as host.

| **Encoded NADP-ME sequence (Name of encoded protein)** | **Vector backbone** | **Production of vector** | **Host *E. coli*** |
|---|---|---|---|
| **C4-NADP-ME** | pET16b | Alvarez *et al.*, 2019 | Rosetta |
| C4-NADP-ME T92F | pET16b | BioCat | Rosetta |
| C4-NADP-ME F140I | pET16b | Alvarez *et al.*, 2019 | Rosetta |
| C4-NADP-ME N142H | pET16b | BioCat | Rosetta |
| C4-NADP-ME N159D | pET16b | BioCat | Rosetta |
| C4-NADP-ME T163R | pET16b | BioCat | Rosetta |
| C4-NADP-ME D164N | pET16b | BioCat | Rosetta |
| C4-NADP-ME V177E | pET16b | SDM | Rosetta |
| C4-NADP-ME G200R | pET16b | BioCat | Rosetta |
| C4-NADP-ME R201Q | pET16b | BioCat | Rosetta |
| C4-NADP-ME D266R | pET16b | BioCat | Rosetta |
| C4-NADP-ME D304H | pET16b | BioCat | Rosetta |
| C4-NADP-ME F317I | pET16b | BioCat | Rosetta |
| C4-NADP-ME M369V | pET16b | BioCat | Rosetta |
| C4-NADP-ME Q377H | pET16b | BioCat | Rosetta |
| C4-NADP-ME I474V | pET16b | BioCat | Rosetta |
| C4-NADP-ME T506N | pET16b | BioCat | Rosetta |
| C4-NADP-ME A521D | pET16b | BioCat | Rosetta |
| C4-NADP-ME Y632F | pET16b | SDM | Arctic Express |
| C4-NADP-ME +15 | pET16b | BioCat | Rosetta |
| C4-NADP-ME N15 | pET16b | BioCat | Rosetta |
| C4-NADP-ME N15 FI | pET16b | SDM | Rosetta |
| C4-NADP-ME +15 FI | pET16b | SDM | Rosetta |
| C4-NADP-ME NnC4 | pET16b | BioCat | Rosetta |
| C4-NADP-ME +DelN | pET16b | BioCat | Rosetta |
| C4-NADP-ME NnC4 G200R | pET16b | SDM | Rosetta |
| C4-NADP-ME N15_G200R | pET16b | SDM | Rosetta |
| **nonC4-NADP-ME** | pET28b | Alvarez *et al.*, 2019 | Rosetta |
| nonC4-NADP-MEΔ15 | pET16b | BioCat | Rosetta |
| nonC4-NADP-MEΔ15_IF | pET16b | SDM | Rosetta |
| nonC4-NADP-MEΔ15_AE | pET16b | SDM | Rosetta |
| nonC4-NADP-MEΔ15_IF_AE | pET16b | SDM | Rosetta |
| nonC4-NADP-MEΔ15_IF_AE_EQ | pET16b | SDM | Rosetta |
| nonC4-NADP-MEΔ15_IF_AE_FL | pET16b | SDM | Rosetta |
| nonC4-NADP-MEΔ15_IF_AE_EQ_FL | pET16b | SDM | Rosetta |
| nonC4-NADP-MEΔ15_13aa mut | pET16b | BioCat | Rosetta |
| nonC4-NADP-MEΔ15_20aa mut | pET16b | BioCat | Rosetta |
| nonC4-NADP-ME I148F | pET28b | SDM | Rosetta |
| nonC4-NADP-ME R171T | pET28b | SDM | Arctic Express |
| nonC4-NADP-ME N172D | pET28b | SDM | Arctic Express |
| nonC4-NADP-ME R208G | pET28b | SDM | Arctic Express |
| nonC4-NADP-ME A347E | pET28b | BioCat | Rosetta |
| **Cyt2** | pET16b | Gibson cloning | Rosetta |
| Cyt2N | pET16b | Gibson cloning | Rosetta |
| Cyt2+DelN | pET16b | SDM | Rosetta |

**Supplementary Table 10. List of primers used in this study.** The table includes the primer name, sequence, and purpose. Lowercase letters indicate modified nucleotides introduced during site-directed mutagenesis. Bold letters denote the regions of Gibson primers that anneal to the target fragment.

| **Name** | **Sequence 5' to3'** | **Purpose** |
|---|---|---|
| FR005 | GATAATGTGGaGGAGCTGC | SDM V177E |
| FR006 | GCAGCTCCtCCACATTATC | SDM V177E |
| SDM C4 Y632F fw | CACTCCCGTCTtCCGCAACTAC | SDM Y632F |
| SDM C4 Y632F rev | GTAGTTGCGGaAGACGGGAGTG | SDM Y632F |
| SDM R163T fw | CAGGAGAcGAACGAGAGG | SDM R171T |
| SDM R163T rev | CCTCTCGTTCgTCTCCTG | SDM R171T |
| SDM N164D fw | CAGGAGAGGgACGAGAGG | SDM N172D |
| SDM N164D rev | CCTCTCGTcCCTCTCCTG | SDM N172D |
| SDM R200G fw | CCATCTTTgGtCAACCACAGG | SDM R208G |
| SDM R200G rev | CCTGTGGTTGaCcAAAGATGG | SDM R208G |
| MH247 | AATTAAGAAGaTCATGAACACCC | SDM F140I |
| MH248 | GGGTGTTCATGAtCTTCTTAATT | SDM F140I |
| MH249 | AAATTAAGAAGtTCATGCACAAC | SDM I148F |
| MH250 | GTTGTGCATGAaCTTCTTAATTT | SDM I148F |
| MH259 | TTAATTTGCTTGaAAAATATAGCAA | SDM A347E |
| MH260 | TTGCTATATTTTtCAAGCAAATTAA | SDM A347E |
| MH267 | GTACTGCTGAAcAAGCATATAACTG | SDM E511Q |
| MH268 | CAGTTATATGCTTgTTCAGCAGTAC | SDM E511Q |
| SDM F544L fw | CCTGGATTaGGCCTCGGTC | SDM F552L |
| SDM F544L rev | GACCGAGGCCtAATCCAGG | SDM F552L |
| FR003 | CATCTTTcGACGACCACAG | SDM G200R |
| FR004 | CTGTGGTCGTCgAAAGATG | SDM G200R |
| sw003 | TAATTTGCTTGcAAAATATAGCA | SDM E347A |
| sw004 | TGCTATATTTTgCAAGCAAATTA | SDM E347A |
| Cyt2 +N-term fwd 1 | GGAGCAGGAGGAGGCGGAGGTGGCCG CGGCC**GCGGGCGGCGGCGTTGAG** | Add nonC4 N-terminal to Cyt2 step 1 |
| Cyt2 +N-term fwd 2 | ATATCGAAGGTCGTCATATGGCCGCGGA GAT**GGAGCAGGAGGAGGCGGAG** | Add nonC4 N-terminal to Cyt2 step 2 + Cloning Zmcyt2 add Gibson overhangs |
| Cyt2 _fwd gibson | ATATCGAAGGTCGTCATATG**GCGGGCGG CGGCGTTGAG** | Clone Cyt2 add Gibson overhangs |
| Cyt2 _rev gibson | GCTTTGTTAGCAGCCGGATC**TTACCGGT AGCTGCGGTAGATGGGG** | Clone Cyt2 add Gibson overhangs |
| Cyt2 _rev full length | TTACCGGTAGCTGCGGTAGATGG | Clone Cyt2 step 1 add N-terminal |
| Cyt2DelN fw | gcaGGCGtCGtCGTTGAGGAC | Exchange Zmcyt2 N-terminal to nonC4 N-terminal, step 1 |
| Cyt2DelN gibson fw 1 | GGAGCAGGAGGAGGCGGAGGTGGCCG CGGCCGCG**GCAGGCGTCGTCGTTGAGG** | Exchange Zmcyt2 N-terminal to nonC4 N-terminal, step 2 |

### Supplementary Table 11. Cryo-EM data acquisition.

| | **nonC4-NADP-ME pH 8.0** | **C4G200R pH 8.0** | **C4G200R pH 4.8** |
|---|---|---|---|
| Magnification | 215,000 | | |
| Voltage (kV) | 300 | | |
| Electron exposure ($e^-/Å^2$) | 50 | | |
| Defocus range (μm) | -0.8 to -2.6 | | |
| Pixel size (Å) | 0.571 | | |
| No. of movies | 3,062 | 4,341 | 876 |
| No. of initial particle images | 447,513 | 393,438 | 140,065 |
| No. of final particle images | 174,901 | 290,820 | 41,462 |
